# Probability of Antibiotic Resistance During Treatment in Stochastic PK/PD-Based Bacterial Model with Distinct Drug and Mutation Modes

**DOI:** 10.64898/2026.06.17.732999

**Authors:** Chimezie Izuazu, Cameron Browne

## Abstract

Mathematical models, e.g. differential equations and stochastic processes, have gained considerable attention for understanding evolution of antibiotic resistance. However, most existing models assume standing genetic variation and do not consider the possibility of random or drug-induced mutation of reference bacterial strains. Therefore, we propose a pharmacokinetics/pharmacodynamics (PK/PD)-based continuous-time Markov chain considering the competition and mutation between sensitive and resistant bacterial within an infected host during treatment. The proposed model is approximated as a generalized birth–death process with immigration, allowing for explicit derivation of the probability resistant population establishes during treatment. Besides capturing the stochasticity of *de novo* emergence of a resistant bacterial strain, we explore the effects of different antibiotic modes of action, horizontal gene transfer, nutrient availability and drug pharmacokinetics on antibiotic resistance. We find that replication-targeting (biostatic) drugs suppress resistance more than death-targeting (biocidal) drugs. Like prior works, we obtain maximized resistance at intermediate drug concentrations, however the consideration of *de novo* mutation magnifies the superiority of higher doses in preventing resistance emergence.

## 1 Introduction

The emergence and fixation of new variants of microbial organisms due to environmental changes have sparked considerable interest among mathematical modelers [1, 2, 3, 4]. The resulting analytical and numerical studies not only validate relevant experimental results [5, 6] but also offer fresh insights into the emergence of drug resistance. In particular, understanding how mutations can rescue of an organism in a deteriorating environment [7, 8] has applications in cancer treatment [9], combination therapy [10], and antibiotic-induced antimicrobial resistance (AMR) mitigation [1]. AMR is one of the major public health threats of the 21st century. Based on published estimates in 2019 [11], 4.95 million people died from AMR-associated illnesses. Particularly, 1.27 million deaths were directly caused by AMR; in fact, drug-resistant infections killed more people than HIV/AIDS (864,000 deaths) or malaria (643,000 deaths). Therefore, understanding AMR evolution is imperative for human sustainability.

Research on AMR has mainly focused on the optimal drug dose and prescription regimen in hospitals to limit the emergence and spread of AMR (see, e.g., [12, 13] and references therein). The question of the optimal drug dose and duration to limit AMR evolution has attracted considerable attention. Pioneering studies, e.g., [14, 15], suggest that “hitting early, hitting hard” limits the *de novo* emergence of AMR. More recent studies (e.g., [16, 17, 18]) challenge this idea based on the result that the probability of AMR emergence is maximized at an intermediate drug concentration. On the one hand, a low drug concentration prevents AMR emergence by allowing the maintenance of the sensitive (wild-type) strain, inhibiting the growth of a resistant strain via competition. On the other hand, a high drug concentration prevents AMR emergence by quickly eradicating the wild-type population, limiting the input of *de novo* resistance mutations and directly limiting the growth of a resistant subpopulation [1]. Therefore, AMR emergence is likely exacerbated at intermediate drug concentrations, where the wild-type population is gradually eradicated, freeing resistance from competition and allowing the resistant subpopulation to grow, i.e., “competitive release” [19]. Depending on the treatment window, which describes the viable drug concentration range and is characterized by the wild-type bacterial clearance and drug toxicity considerations, a low or high drug dose may be optimal to limit population rescue via AMR emergence [1].

Several studies (e.g., [20, 21, 22, 23]) deterministically modeled the pharmacokinetics (PK)/pharmacodynamics (PD)–based dynamics of sensitive and resistant cells *in vitro* or *in vivo* to describe how a resistant strain can grow and rescue the bacterial population. This deterministic description is mainly justified when a substantial resistant subpopulation exists before treatment. However, in several common situations, the resistant bacterial subpopulation is initiated from none or few resistant cells, and mutations or gene transfers conferring resistance rarely occur. This early phase of resistance emergence is a stochastic process. Theoretical studies of the stochastic effect on AMR emergence have often required various simplifying assumptions to make progress [1, 8, 12, 18, 24]. One method has been to exploit the orders of magnitude of the population size difference between drug sensitive and resistant strains initially by assuming the former has independent deterministic dynamics whereas the latter is governed by a stochastic process [1, 4].

Naturally, under the deteriorating environmental conditions induced by antibiotics, bacterial organisms seek means to adapt through mutation and selection [25]. However, most existing numerical and analytical solutions for resistance survival probabilities under various treatment conditions assume standing genetic variation (SGV) or that a new bacterial strain is introduced from an external source but do not consider the possibility of pure mutations of the sensitive wild-type bacterial strain (see, e.g., [1, 2, 3, 4]). In addition, the effects of distinct mutation mechanisms, such as horizontal gene transfer (HGT) and drug-induced mutations, along with explicit PK dynamics, have been relatively under-explored. Such considerations could be important to further characterize the drug concentration maximizing the probability of AMR emergence, the size of the resistant subpopulation within the host and, in the case of commensal bacteria, how long a treated host carries and sheds resistance after treatment. Moreover, a stochastic description of AMR evolution requires that we specify whether the drug inhibits cell division or kills cells (biostatic or biocidal) as an important determinant of the survival probability of emergent resistant cells [1]. Here, we focus on characterizing the drug concentration maximizing the probability of AMR emergence under the different drug types, explicit mutation modes, and PK/PD dynamics.

In this study, we propose a PD-based continuous-time Markov chain (CTMC) model that considers the emergence of *de novo* bacterial strain by pure mutations of the wild-type strain and incorporate PK to provide in-depth insights into the mechanisms of antibiotic-induced AMR. The proposed model can capture the stochasticity of both *de novo* emergence of a bacterial strain and HGT via conjugation, thereby offering insights into the effects of these two sources of AMR on the antibiotic-induced rescue or eradication of a bacterial population. Under mean-field approximation of the wild-type population, the proposed model is tractable as a generalized birth–death process (GBDP), which is time inhomogeneous [26]. Particularly, under *de novo* mutation, our model is a GBDP with immigration (GBDP-I) [27]. Although similar frameworks have been recently analyzed [1, 10, 28], they either do not consider explicit *de novo* mutation and PK [1], neglect resource competition, HGT, and directed mutation [10], or do not consider drug mode of action [28]. This study aims to illuminate the combined effects of these factors on AMR evolution.

The remainder of this article is organized as follows. In Section 2, we describe the proposed model as well as present relevant definitions and standard results for GBDP and GBDP-I. Theoretical analysis of the proposed model is presented in Section 3. In Section 4, we consider a concrete scenario and present analytical results of survival/establishment probability of resistant bacterial cells and wild-type clearance. Moreover, we present simulations of various dose-dependent mutations and analyze the effect of nutrient availability on antibiotic resistance and susceptibility, answering a recently posed question [29]. Further, we incorporate PK into the proposed model for in-depth insights into the mechanism of antibiotic resistance. Finally, we discuss our results, conclude this study, and suggest future research directions in Section 5.

## 2 Model

Let {*w*_*t*_}_*t*≥0_ and {*m*_*t*_}_*t*≥0_ be stochastic processes for the numbers of wild-type (sensitive) and mutant (resistant) cells, respectively. Thus, {*N*_*t*_ := *w*_*t*_ + *m*_*t*_}_*t*≥0_ is the stochastic process for the total bacterial population. In addition, let *c*(*t*; *c*_0_) := *c* be the drug concentration at time *t*, which can differ from the initial drug concentration, *c*_0_, when PK is considered. We model the bacterial population dynamics as the following multi-type CTMC:

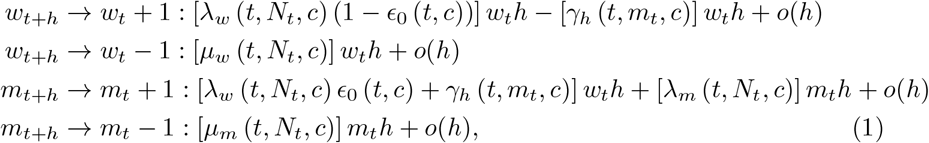

where *w*_*t*+*h*_ → *w*_*t*_ ± 1 (resp. *m*_*t*+*h*_ → *m*_*t*_ *±* 1) means the instantaneous event that a single cell enters or leaves the wild-type (resp. mutant) strain compartment, *λ*_*i*_(.) and *µ*_*i*_(.) denote the bacterial birth and death rates, respectively (*i* ∈ {*w, m*}), *ϵ*_0_(.) denotes the *de novo* mutation rate, and *γ*_*h*_(.) denotes the HGT rate. The explicit dependence of these rates on time and drug concentration reflects the effects of external environmental changes and antibiotics on the population dynamics, respectively [1]. We assume *γ*_*h*_(*t, m*_*t*_, *c*) ≤ *λ*_*w*_ (*t, N*_*t*_, *c*) (1 − *ϵ*_0_(*t, c*)), where *ϵ*_0_(*t, c*) denotes the *de novo* mutation rate and *γ*_*h*_(*t, m*_*t*_, *c*) denotes the HGT rate.

**Table 1.** Summary of the different birth and death events in the population, along with the associated rates per individual, and the total rate for any type of event in the population. The symbol ∅ represents removal of an individual from the population (i.e., death).

| Transition | per capita rate | change of $(w_t, m_t)$ | Total event rate |
| --- | --- | --- | --- |
| $w \rightarrow w + w$ | $\lambda_w(t, N_t, c) (1 - \epsilon_0(t, c)) - \gamma_h(t, m_t, c)$ | $(1, 0)$ | $[\lambda_w(t, N_t, c) (1 - \epsilon_0(t, c)) - \gamma_h(t, m_t, c)] w_t$ |
| $w \rightarrow \emptyset$ | $\mu_w(t, N_t, c)$ | $(-1, 0)$ | $[\mu_w(t, N_t, c)] w_t$ |
| $w \rightarrow w + m$ | $\lambda_w(t, N_t, c) \epsilon_0(t, c) + \gamma_h(t, m_t, c)$ | $(0, 1)$ | $[\lambda_w(t, N_t, c) \epsilon_0(t, c) + \gamma_h(t, m_t, c)] w_t$ |
| $m \rightarrow m + m$ | $\lambda_m(t, N_t, c)$ | $(0, 1)$ | $[\lambda_m(t, N_t, c)] m_t$ |
| $m \rightarrow \emptyset$ | $\mu_m(t, N_t, c)$ | $(0, -1)$ | $[\mu_m(t, N_t, c)] m_t$ |

In general, we consider a dose-dependent mutation rate *ϵ*_0_(*t, c*) and massaction-type HGT 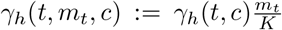 as the two pathways to generate resistant cells from wild-type cells. These pathways decrease and increase wildtype and mutant cell replications, respectively, thereby reshaping their fitness costs and benefits. Unlike spontaneous or constant mutation, the literature on dose-dependent mutation models is inadequate [25, 30]. Nevertheless, under antibiotics-induced environmental changes, bacterial organisms seek means to adapt via mutation and selection [25], rationalizing a dose-dependent mutation model. Moreover, defining *ϵ*_0_(*t, c*) as a constant function recovers the more prominent spontaneous mutation. Meanwhile, we consider a mass action HGT function [31], such that the frequency of mutant cells that emerge via HGT is proportional to the relative frequencies of both strains and is controlled by *γ*_*h*_(*t, c*), which depends on the environmental conditions.

We also assume total resistance to treatment, i.e., *λ*_*m*_ (*t, N*_*t*_, *c*) = *λ*_*m*_ (*t, N*_*t*_, 0) and *µ*_*m*_ (*t, N*_*t*_, *c*) = *µ*_*m*_ (*t, N*_*t*_, 0), ∀*c >* 0, *t* ≥ 0, although our analysis can be modified for partial resistance, i.e., resistance that merely increases the minimum inhibitory concentration (MIC); we simulate this modification in the Supplementary Information (SI; Section SI2). Further, we assume that either the resistant bacterial subpopulation does not exist or is relatively minute at the beginning of treatment such that its effect on the rates and dynamics of the sensitive bacterial strain is negligible.

Because the sensitive bacterial subpopulation is relatively large and its dynamics are dominated by antibiotic-induced decay, following standard practice in ecological modeling [1], we ignore stochastic fluctuations in the subpopulation dynamics by replacing the stochastic variable, *w*_*t*_, with its expected value, *w*(*t*) := E[*w*_*t*_], resulting in the following deterministic equation:

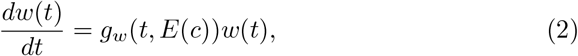

where *g*_*w*_(*t, E*(*c*)) := [*λ*_*w*_(*t, w*(*t*), *c*) (1 − *ϵ*_0_(*t, c*)) − *µ*_*w*_(*t, w*(*t*), *c*)] is the drug-mediated growth rate, with *E*(*c*) as the drug effect function. Notably, we approximate the rate functions as independent of the resistant subpopulation because its dynamics are stochastic and its size is negligible.

In this study, we consider the following effect function [32]:

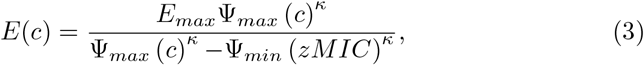

where *κ* denotes the Hill coefficient, *E*_*max*_ denotes the maximum antibiotic-mediated death rate, Ψ_*max*_ denotes the death rate in the absence of antibiotics, *zMIC* denotes the PD MIC, and Ψ_*min*_ = Ψ_*max*_ − *E*_*max*_ denotes the minimum growth rate at high antibiotic concentration. Moreover, we consider two modes of action of drugs—biostatic and biocidal—where a drug is biostatic if it inhibits birth rate independent of death rate or biocidal if it promotes death rate independent of birth rate [1].

## 3 Theoretical Analysis

Here, we present theoretical analysis of the proposed model and some biological interpretations. After solving the deterministic equation for *w*(*t*; *c*), based on the assumption that the effect of the resistant bacterial subpopulation is negligible, the following approximations are made: *λ*_*w*_(*t, c*) := *λ*_*w*_ (*t, w*(*t*), *c*) ≈ *λ*_*w*_ (*t, N*_*t*_, *c*); *λ*_*m*_(*t, c*) := *λ*_*m*_ (*t, w*(*t*), *c*) ≈ *λ*_*m*_ (*t, N*_*t*_, *c*); *µ*_*m*_(*t, c*) := *µ*_*m*_ (*t, w*(*t*), *c*) ≈ *µ*_*m*_ (*t, N*_*t*_, *c*); and *µ*_*w*_(*t, c*) := *µ*_*w*_ (*t, w*(*t*), *c*) ≈ *µ*_*w*_ (*t, N*_*t*_, *c*). Moreover, 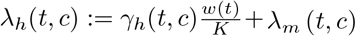. Thus, we obtain the following GBDP-I for the resistant bacterial sub-population:

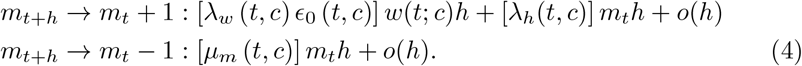

Let 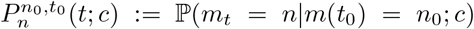, i.e., the probability that the treatment-induced number of resistant cells at time *t* is *n* given that the number of resistant cells at time *t*_0_ (initial time) is *n*_0_. Based on first principles, we obtain the following difference–differential equation:

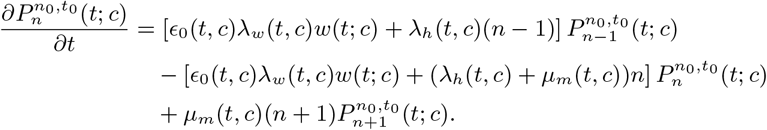

In addition, let

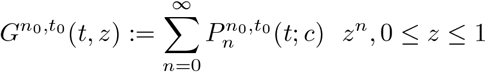

be the generating function, which satisfies the following partial differential equation (PDE):

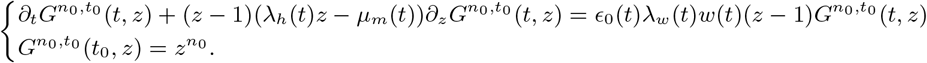

Notably, we suppress the explicit dependence on drug concentration, *c*; however, this is not a limitation because the following generic calculations can be performed without suppressing *c*.

Following Allen [33], we use the method of characteristics to solve the above PDE, which is quasillinear (see Appendix B).

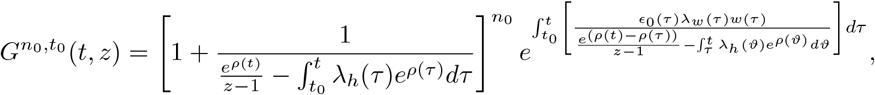

where *ρ*(*t*) := [*µ*_*m*_(*t*) *λ*_*h*_(*t*)] *dt*.

Moreover, observe that

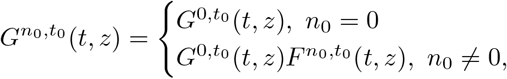

where

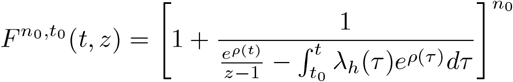

is the generating function of GBDP. Therefore, we adopt the following procedure in determining the transition probability, 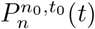 [27]:

1. calculate the GBDP’s transition probability, 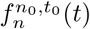;
2. calculate the GBDP-I’s zero-state transition probability, 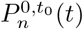;
3. calculate the GBDP-I’s transition probability, 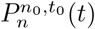, as the convolution of 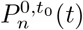 and 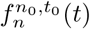.

Based on this procedure, we determine the GBDP-I’s transition probability, 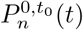. To this end, we first recall the following result for GBDP.

### Proposition 1

(Bailey et al., [34]). *For t* ≥ *t*_0_ *and* 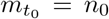, *the GBDP*’*s transition probability*, 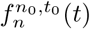, *is given by*

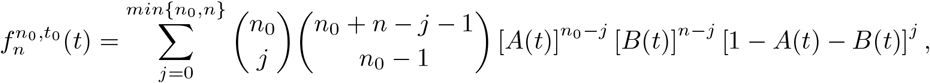

*where*

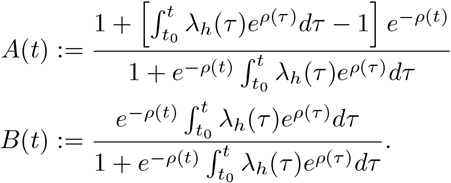

*In particular*,

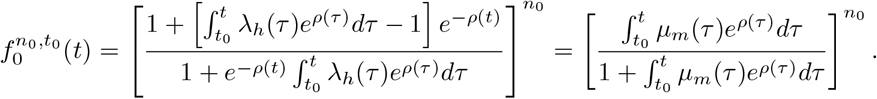

Next, we calculate 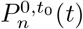 based on the complete Bell polynomials employed by Virginia et al. [27].

### Proposition 2

*For t* ≥ *t*_0_, *the GBDP-I*’*s zero-state transition probability*, 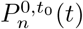, *is given by*

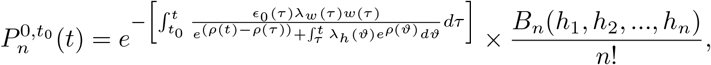

*where B*_*n*_’*s are the complete Bell polynomials recursively given by* 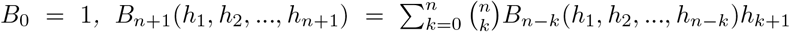 with 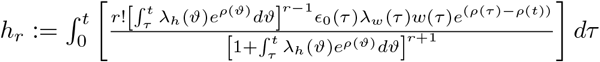

*In particular, the extinction probability of de novo mutation is given by*

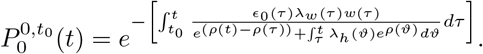

**Proof** See Appendix C

### Proposition 3

*For t* ≥ *t*_0_ *and n*_0_≠ 0, *the transition probability* 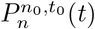 *of GBDP-I is given by*

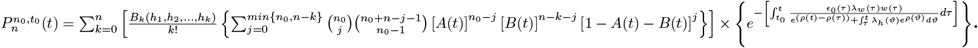

*In particular, for n* = 0, *we have*

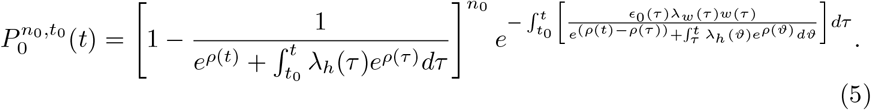

**Proof** Follows directly from Propositions 1 and 2 and the above outlined procedure.

Further, the following identity can be realized via integration by substitution.

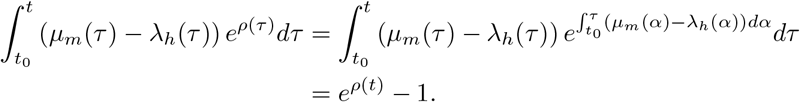

Thus,

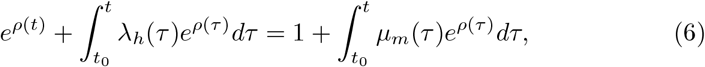

which yields

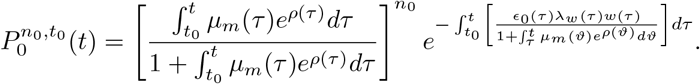

Let 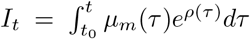 and 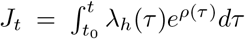. Thus, the extinction probability of resistant bacterial cells can be expressed as follows:

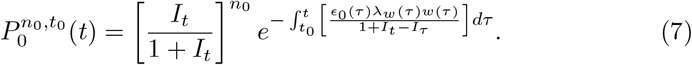

Consequently, without SGV (i.e., *n*_0_ = 0), we obtain the extinction probability of *de novo* resistant bacterial cells as follows:

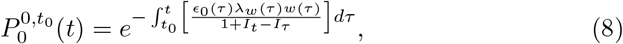

which corresponds to the nonoccurrence probability of a time-inhomogeneous Poisson process. This validates Teemu et al.’s [35] model of the acquisition of rescue mutants by an inhomogeneous Poisson process. The integrand, 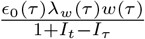, can be interpreted as the rate of generating resistant bacterial cells at time *τ* ∈ [*t*_0_, *t*], with *ϵ*_0_(*τ*), *λ*_*w*_(*τ*), and *w*(*τ*) denoting the per ca-pita *de novo* mutation rate, per capita birth rate of sensitive bacteria, and sensitive bacterial population at time *τ*, respectively, and 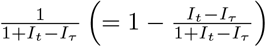 quantifying the probability of a *de novo* resistant bacterial cell that appeared at time *τ* to survive at time *t*. Therefore, the survival probability of the resistant bacteria strain at time *T* can be expressed as follows:

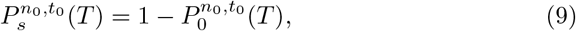

which is comparable to Teemu et al.’s [35] probability of an evolutionary rescue by *de novo* mutation and Ernesto et al.’s [28] expression of the probability of a single-resistant cell after the treatment commencement. In addition, we can express the establishment probability of the resistant bacteria as follows, when the corresponding limit exists^1^:

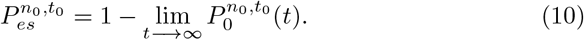

We also use the following notations: 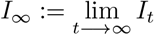 and 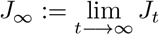.

Further, on the one hand, if *µ*_*m*_ ≡ 0, then 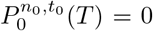, i.e., there is almost surely no chance of extinction; on the other hand, if *µ*_*m*_(*t*) = 0, for some *t* ≥ 0, then 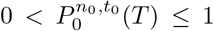, i.e., there is a chance of extinction. Subsequently, unless stated otherwise, we assume a non-zero death rate.

## 4 Results

As a concrete scenario, we consider logistic birth rates, 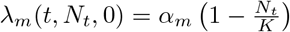 and 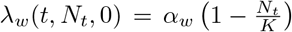, and constant death rates, *µ*_*m*_(*t, N*_*t*_, 0) = *µ*_*m*_ and *µ*_*w*_(*t, N*_*t*_, 0) = *µ*_*w*_, where *µ*_*m*_, *µ*_*w*_, *α*_*m*_, and *α*_*w*_ are positive constants. We also assume *N*_*t*_ ≤ *K*, which ensures that the natural birth and death rates of both strains are non-negative.

### 4.0.1 Biocidal Treatment

For biocidal treatment, which promotes death rate independent of birth rate [1], the drug-mediated growth rate is given by

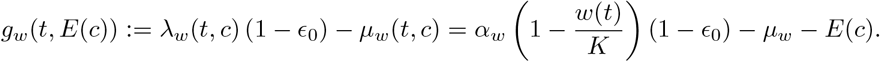

Consequently, we obtain the closed-form solution of the wild-type dynamics as follows:

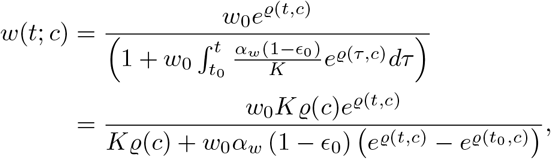

where ϱ(*c*) := [*α*_*w*_ (1 − *ϵ*_0_) − *µ*_*w*_ − *E*] and ϱ(*t, c*) := ϱ(*c*)*t*.

### 4.0.2 Biostatic Treatment

For biostatic treatment, which inhibits birth rate independent of death rate [1], the drug-mediated growth rate is given by^2^

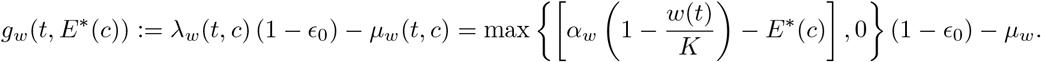

Notably, if 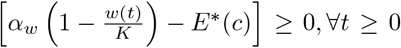 ^3^, which is guaranteed at low drug concentrations, then the growth rate simplifies as follows:

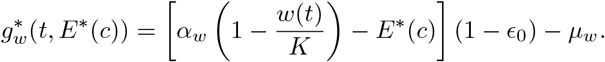

Consequently, we obtain the closed-form solution of the wild-type dynamics as follows:

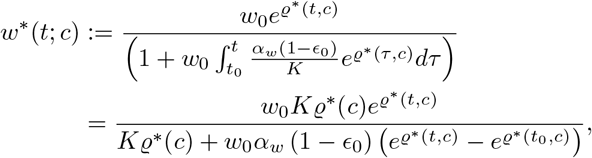

where ϱ^∗^(*c*) := [*α*_*w*_ (1 − *ϵ*_0_) − *µ*_*w*_ − (1 − *ϵ*_0_) *E*^∗^] and ϱ^∗^(*t, c*) := ϱ^∗^(*c*)*t*.

Subsequently, to capture a realistic viable treatment strategy, we assume that the drug-mediated intrinsic growth rate of the sensitive bacteria strain, ϱ^(∗)^(*c*), is non-positive. Table 2 summarizes the wild-type dynamics for the two drug modes of action.

**Table 2.** Wild-type dynamics for PD biocidal vs. biostatic treatment.

| Mode of Action | Biocidal | Biostatic |
| --- | --- | --- |
| Intrinsic growth rate | $\varrho(c) = [\alpha_w (1 - \epsilon_0) - \mu_w - E]$ | $\varrho^*(c) = [\alpha_w (1 - \epsilon_0) - \mu_w - (1 - \epsilon_0) E^*]$ |
| Expected wild-type dynamics | $w(t; c) = \frac{w_0 e^{\varrho(c)t}}{\left( 1 + w_0 \int_{t_0}^t \frac{\alpha_w (1 - \epsilon_0)}{K} e^{\varrho(c)\tau} d\tau \right)}$ | $w^*(t; c) = \frac{w_0 e^{\varrho^*(c)t}}{\left( 1 + w_0 \int_{t_0}^t \frac{\alpha_w (1 - \epsilon_0)}{K} e^{\varrho^*(c)\tau} d\tau \right)}$ |
| Constraint | – | $0 \leq E^* \leq \alpha_w \left( 1 - \frac{w(t)}{K} \right)$ |

Notably, under biostatic treatment, we set 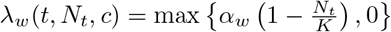 because birth rate cannot be less than zero. However, this is not enforced under the biocidal treatment because death rate can, in principle, increase without bound as the antibiotic concentration increases [1]. Table 3 summarizes the parameters used in this study and their literature values.

**Table 3.** Ciprofloxacin-adapted PD parameters for bacterial evolution. Besides the referenced parameter values, we choose *α*_*w*_ *& µ*_*w*_ such that *α*_*w*_ −*µ*_*w*_ = Ψ_*max*_; *α*_*m*_ *& µ*_*m*_ such that the wild-type strain has a fitness advantage in the absence of antibiotics; and *w*_0_ & *m*_0_ such that the bacterial population is almost saturated at the carrying capacity Population size

| Parameter | Meaning | Value (unit) | Source |
| --- | --- | --- | --- |
| $\kappa$ | Hill coefficient | 1.1 | [32] |
| $\Psi_{min}$ | Minimum growth rate at high antibiotic concentration | $-6.5 \text{ (h}^{-1}\text{)}$ | [32] |
| $E_{max}$ | Maximum antibiotics-mediated death rate | $7.38 \text{ (h}^{-1}\text{)}$ | [32] |
| $zMIC$ | MIC for sensitive wild-type bacteria | $0.017(\mu\text{gml}^{-1})$ | [32] |
| $\Psi_{max}$ | Growth rate in the absence of antibiotics | $0.88 \text{ (h}^{-1}\text{)}$ | [32] |
| $nMIC$ | MIC for mutant bacteria | $n \times zMIC, n \in [0, \infty] (\mu\text{gml}^{-1})$ | [1] |
| $\mu_m$ | Natural mutant death rate | $2.5 \text{ (h}^{-1}\text{)}$ | |
| $\mu_w$ | Natural wild-type death rate | $2.5 \text{ (h}^{-1}\text{)}$ | |
| $\alpha_m$ | Maximum natural mutant birth rate | $2.88 \text{ (h}^{-1}\text{)}$ | |
| $\alpha_w$ | Maximum natural wild-type birth rate | $3.38 \text{ (h}^{-1}\text{)}$ | |
| $K$ | Carrying capacity | Variable | |
| $T$ | Treatment duration | $\leq 72 \text{ h}$ | |
| $w_0$ | Initial wild-type size | $0.99 \times K$ | |
| $m_0$ | Initial resistant size | Minute | |

Figures 1 and 2 show sample paths for the original multi-type CTMC (1) and GBDP-I approximation (4), respectively, with logistic birth rate, constant death rate, and constant mutation rate. For comparison, we present the corresponding resistant bacterial subpopulation dynamics of the original CTMC model (hereinafter, resistant sub-CTMC) as insets. Clearly, the GBDP-I and resistant sub-CTMC follow the same qualitative trend, particularly at the initial treatment phase, validating the GBDP-I approximation. Notably, the numerical simulations are based on the *τ* -leaping approximate accelerated stochastic simulation algorithm [36].

**Figure 1.**
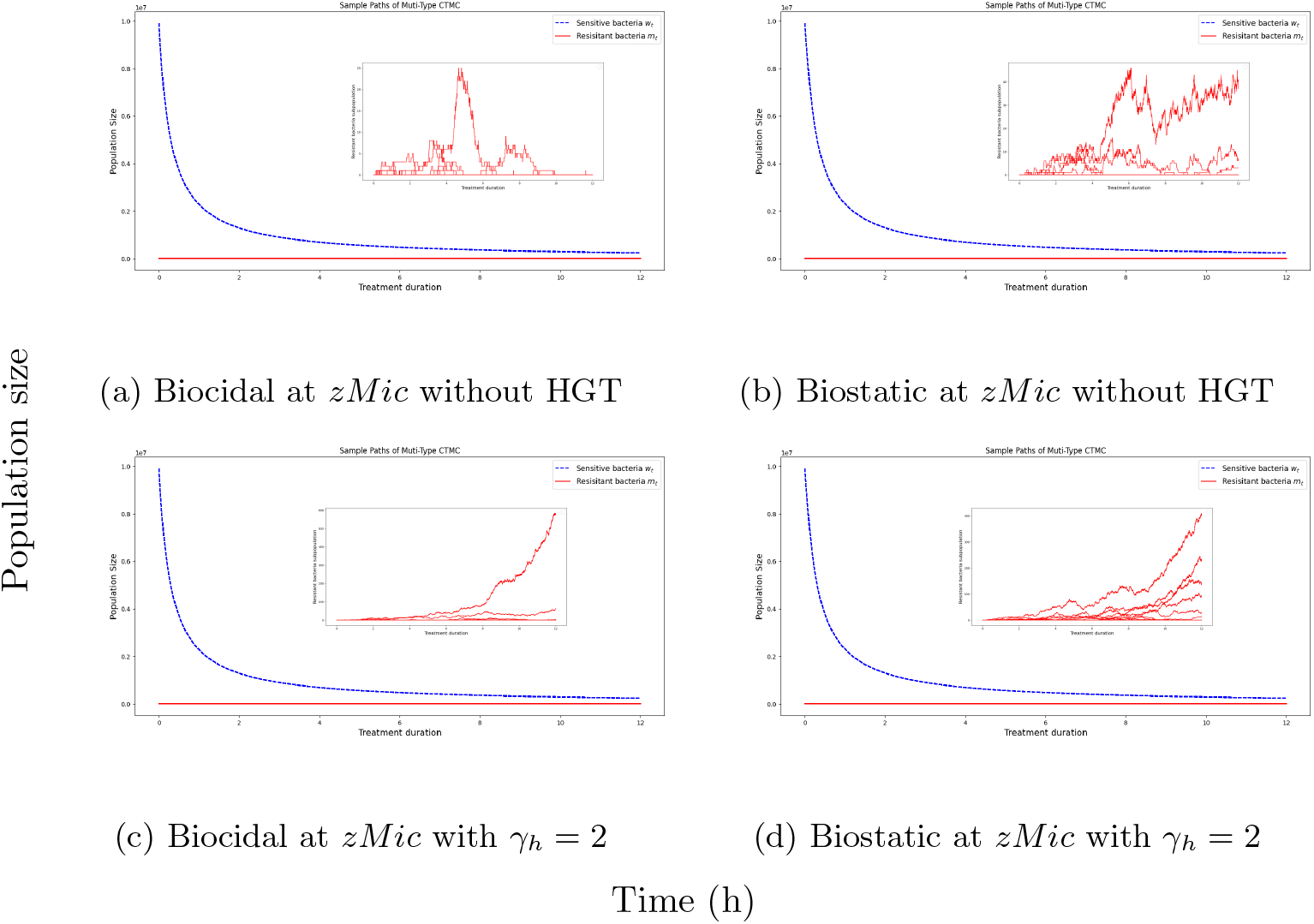
Population dynamics of sensitive (blue) and resistant (red) strains under logistic growth conditions within 12-*h* PD-based treatment, with *K* = 10^7^, *ϵ*_0_ = 10^−8^ and *m*_0_ = 0 (*de novo*) for 100 sample paths; the update terminates if the total change rate or count of the wild-type subpopulation is zero or below 1, respectively. The insets are magnifications of the resistant subpopulation dynamics, which is mainly governed by stochastic drift.

**Figure 2.**
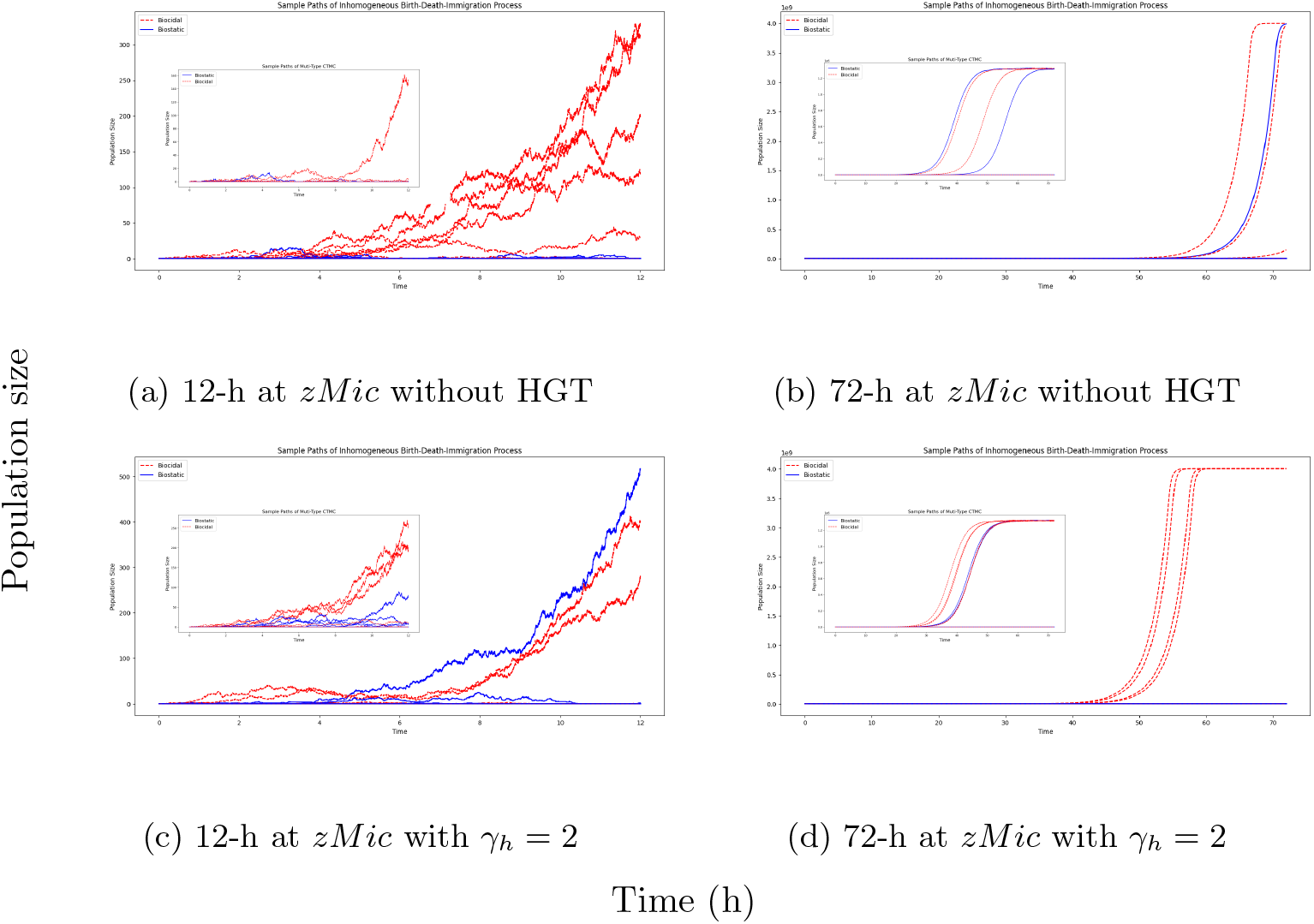
Short- and long-term population dynamics of the resistant bacterial strain under logistic growth conditions at constant drug concentrations for a biocidal drug (ciprofloxacin; red) and its biostatic analog (blue), with *K* = 10^7^, *ϵ*_0_ = 10^−8^ and *m*_0_ = 0 (*de novo*) for 100 sample paths. The insets indicate resistant sub-CTMC for comparison.

### 4.1 PD-Based Dynamics

#### 4.1.1 Survival Chance of Resistant Strain

At time *T* ≥ 0, the extinction probabilities of the resistant strain under biocidal and biostatic treatment conditions are, respectively, expressed as follows:

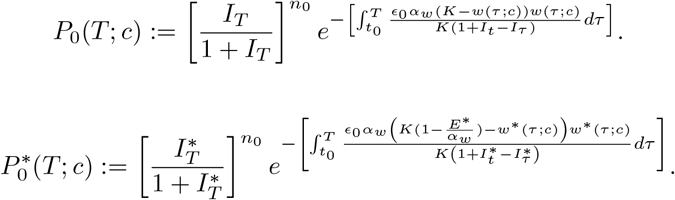

Further, in a finite state space, the resistant strain either goes extinct (zero cells) or survives (one or more cells). Therefore, the survival probabilities of the resistant strain at time 0 ≤ *T <* ∞ for the biocidal and biostatic treatments are, respectively, given by

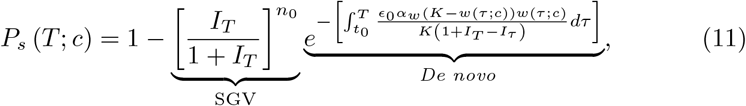

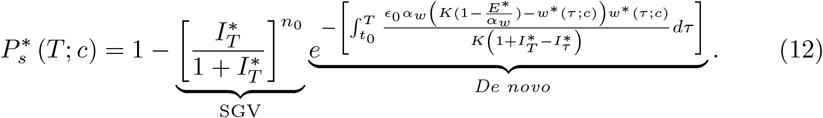

Letting *T* → *∞*, we obtain the corresponding resistant bacterial establishment probabilities:

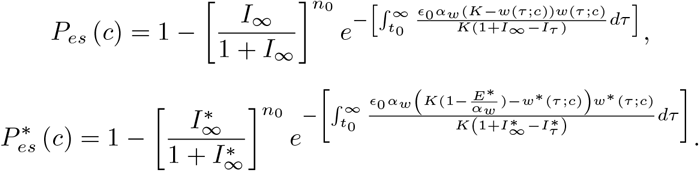

In general, the establishment probabilities seem analytically intractable. However, under the special condition that *α*_*m*_ = *α*_*w*_ (1 − *ϵ*_0_) + *γ*_*h*_, they can be simplified as follows^4^:

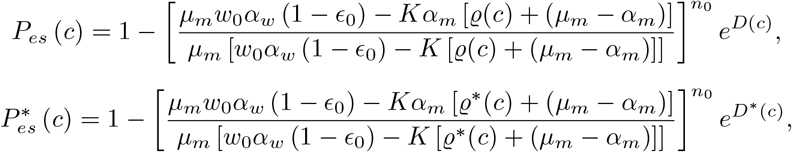

Where 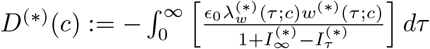, which quantifies the amount of *de novo* mutations, converges for the relevant parameter values. Further, neglecting resistance via *de novo* mutations, i.e., *D*^(∗)^(*c*) = 0, and assuming that *n*_0_ = 1, we obtain the following simplifications of the establishment probabilities of the resistant cells:

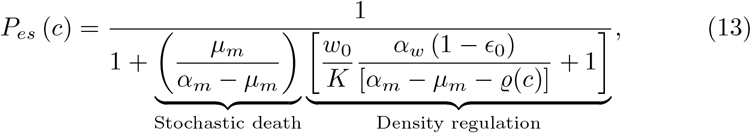

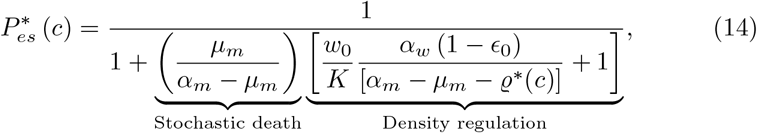

These concise equations, which are comparable to Czuppon et al.’s [1] result (i.e., Eq. (3)), elucidate the factors influencing the resistant establishment probability other than mutation. They encapsulate, in simple forms, the chance that a single resistant cell survives treatment under SGV with an initial sensitive subpopulation of size *w*_0_, which decreases under the action of treatment. The establishment probability is large when the factor in the denominator is small, depending on two processes—stochastic variability and competition via density regulation. For stochastic variability, the stochastic death of resistant cells threatens the survival of resistant cells. This is translated in mathematical terms by the contribution of the resistant death rate to the maximum growth rate (the term 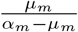). Further, this term increases with *µ*_*m*_, thereby reducing the establishment probability. Of course, if the resistant cells do not die, *µ*_*m*_ = 0, the establishment probability is 1, i.e., the resistant strain will almost surely be established. The density regulation is mediated by the term 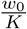, which is scaled by the factor 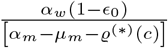, indicating that competition is alleviated as the drug-induced growth rate of the sensitive strain reduces. In addition, if *E* ≡ *E*^∗^, then 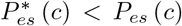, indicating the superiority of the biostatic analog for long-term treatment. Notably, analogous formulas can be obtained for a finite treatment duration, say *T* . Further, letting *K* approach infinity (i.e., infinite resources), we obtain

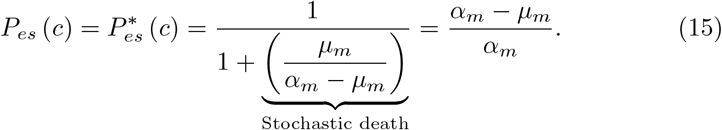

This equality of biostatic- and biocidal-based establishment chances under infinite resources, perhaps, explains the meta-analysis results of no difference in treatment success between both drug modes for severe infections [37, 38].

As opposed to the above special case, we now derive some potential differences between biostatic and biocidal treatment. First, the following analytical result (Theorem 1) indicates a faster clearance of the wild-type strain with a biocidal drug, consistent with previous studies (see, e.g., [1, 39], and references therein).

##### Theorem 1

*Suppose E* ≡ *E*^∗^ *(i*.*e*., *equality of biostatic and biocidal drug effects). Then, w*(*t*; *c*) ≤ *w*^∗^(*t*; *c*), ∀*t* ≥ 0.

**Proof** Let 1 ≥ *ϵ*_0_ *>* 0 and ∞ *> T >* 0 be fixed. Assume *E* ≡ *E*^∗^, we have that 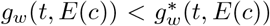. Therefore, by the basic comparison theorem of ODEs,

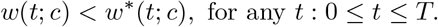

Further, because *T* is arbitrary, we have that

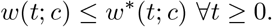

Moreover, we consider existing literature parameters [1, 32] (Table 3) to validate this result numerically. Figure 3 shows the plots of the logarithm of wild-type count against treatment duration. The numerical values indicate that a biocidal drug clears bacterial cells faster than its biostatic analog, consistent with our theoretical result. Moreover, the efficacy of a biocidal drug in clearing wild-type bacterial cells is more pronounced for a shorter treatment duration at a higher drug concentration (Figure 3(a)). These findings seem intuitive, as the entire biocidal effect directly eliminates sensitive bacterial cells whereas a portion of the biostatic effect dampens the mutation effect due to the explicit mutation rate.

**Figure 3.**
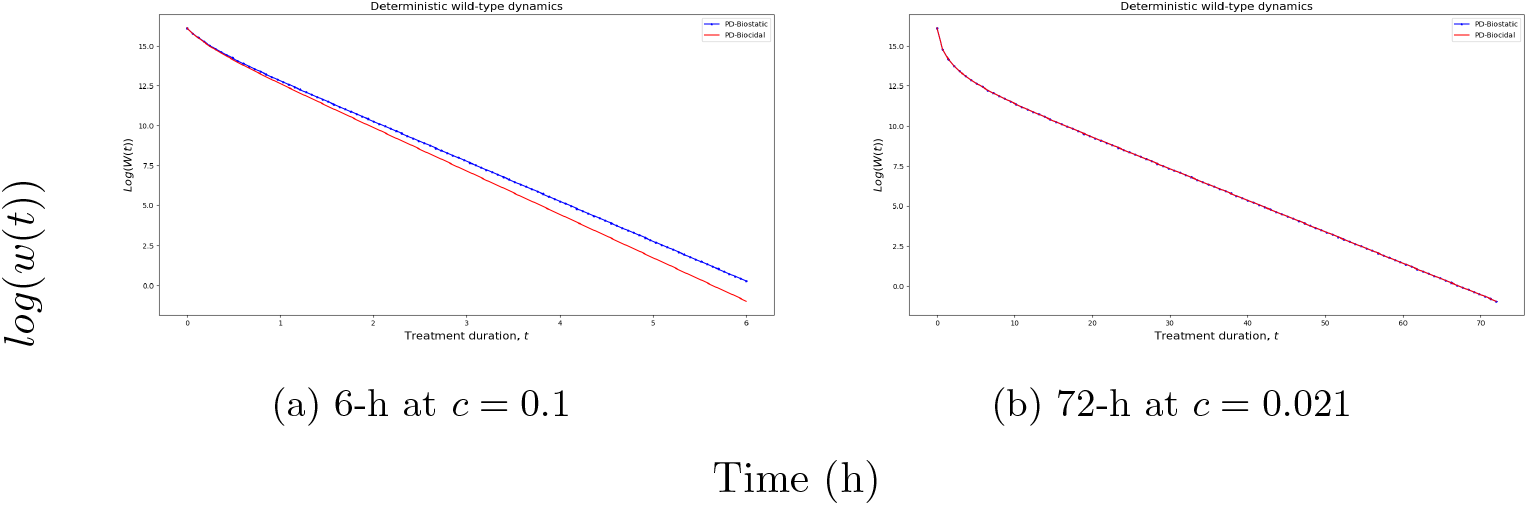
Wild-type clearance rate for PD biocidal vs. biostatic treatment with *K* = 10^7^ and *ϵ*_0_ = 10^−7^.

Nevertheless, Theorem 2 highlights the superiority of a biostatic drug in suppressing the survival probability of resistant cells when the mutation rate is sufficiently small.

##### Theorem 2

*Assuming E* ≡ *E*^∗^. *Suppose T <* ∞. *Then*, ∃ Δ *>* 0 *such that* 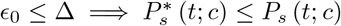 *for any t* ∈ [0, *T*]. *Moreover, we can estimate a suitable* Δ *as follows:*

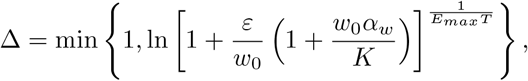

*where ε >* 0 *is a chosen bound on the maximal wild-type density difference, w*^∗^(*t*; *c*) − *w*(*t*; *c*) *< ε*.

**Proof** See Appendix D

Figure 4 shows plots of the survival probability of *de novo* resistant cells against drug concentration under biostatic and biocidal treatment conditions at various HGT rates and a fixed mutation rate. Notably, the survival probability under both treatment conditions peaks at sub-MIC levels. In addition, the numerical results indicate that the biostatic drug has a higher potency in decreasing the survival probability of *de novo* resistant bacteria. This seems intuitive because a biostatic drug inhibits cell replication, thereby suppressing the emergence of *de novo* bacterial cells. Overall, the biocidal-based survival chance does not exceed the biostatic-based survival chance, consistent with Theorem 2. These results, combined with insights from Figure 3, suggests that a biocidal drug is suitable for short-duration treatment at higher drug concentrations (considerably greater than *zMIC*) whereas its biostatic analog is suitable for long-duration treatment at lower drug concentrations.

**Figure 4.**
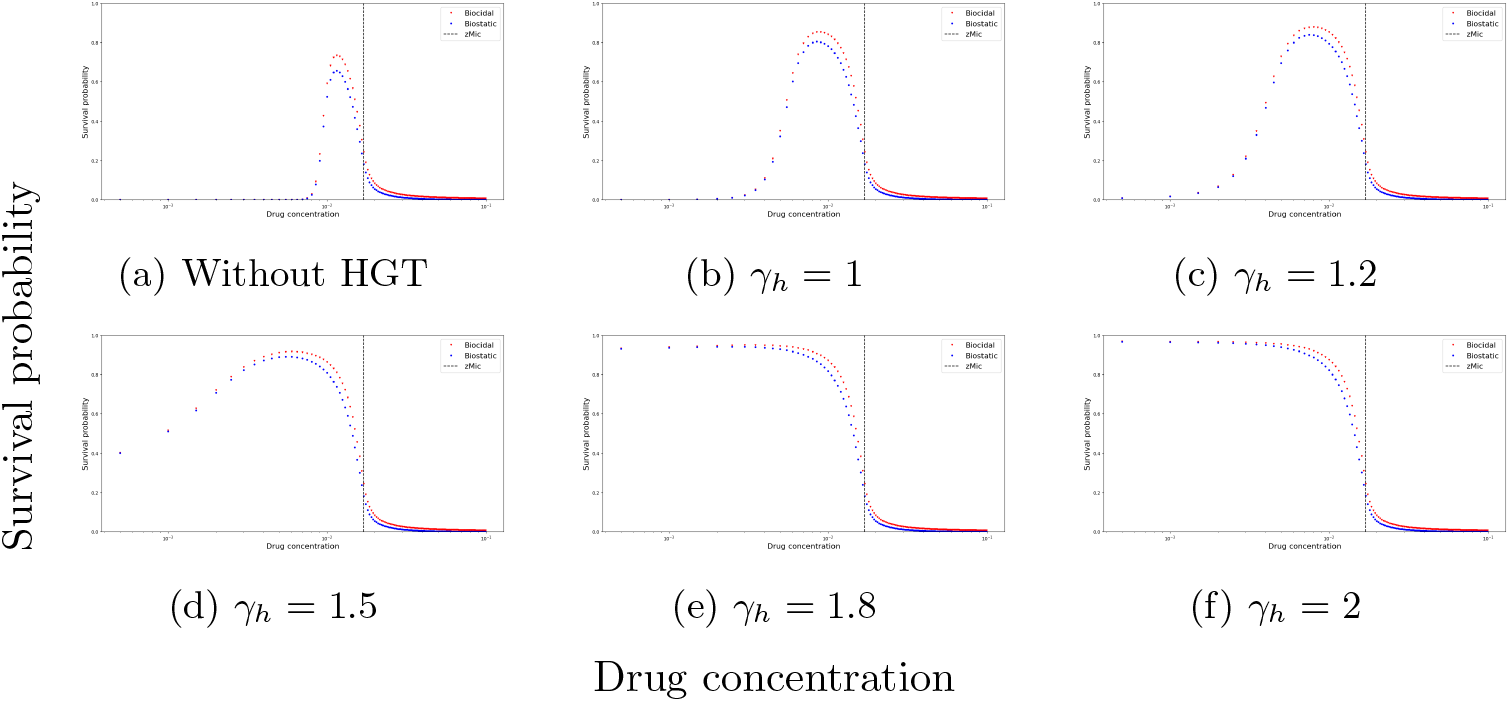
Survival probability of *de novo* resistant bacteria against drug concentration for 72-h PD-based biostatic vs. biocidal treatments with *K* = 10^7^ and *ϵ*_0_ = 10^−8^.

The above results highlight the superiority of biostatic treatment in suppressing *de novo* resistant bacterial establishment. We infer that the explicit mutation dampens the biostatic drug efficacy, as some of the effect results in fewer mutants being produced rather than directly reducing the sensitive sub-population. Another factor, independent of the mutation rate, is that the birth rate must be non-negative, so drug effects that reduce the birth rate to less than zero are not considered, as emphasized by Czuppon et al. [1]. However, the first factor of explicit mutation dampening the biostatic drug effect is offset by fewer mutants being born due to the drug effect, thereby reducing the survival probability. In addition, as noted by Czuppon et al. [1], the slower clearing of the wild-type cells by biostatic treatment than by biocidal treatment allows less competitive release, thereby further reducing the survival probability.

Figure 5 shows plots of survival probability against drug concentration under biocidal and biostatic treatment conditions at various HGT rates considering pre-existing single resistant cells (SGV) besides *de novo* mutations. Consistent with Theorem 2, the biostatic drug has a higher potency in decreasing the survival probability, particularly at sub-MIC levels without HGT. Under SGV, the survival probability tends to converge to a non-zero value as the drug concentration increases, unlike the consideration of only the *de novo* resistant strain in which “almost sure” suppression can be guaranteed at high drug concentrations. This indicates that aggressive treatments under SGV enhance the selective advantage of resistant cells, attributable to the well-known phenomenon of competitive release in which treatment-induced selection pressure not only leads to the decay of sensitive cells but also facilitates the growth of pre-existing resistant cells. From Figures 4 and 5, it is obvious that HGT is a major driver of resistant bacterial establishment for treatment at sub-MIC levels. Particularly, high HGT rates shift the survival probability dynamics from non-monotonic to a monotone decreasing trend with increasing drug concentration, indicating that HGT facilitates more sensitive competitive release, particularly at sub-MIC levels. Although treatments at sub-MIC levels reduce the sensitive bacterial population, they allow cell replication, which can facilitate antibiotic-induced population rescue via the combined effect of *de novo* mutation and HGT. This population rescue is further strengthened under SGV, and longer-duration treatments at sub-MIC levels will “almost surely” guarantee sensitive bacterial establishment. Notably, the pronounced disadvantage of treatments at sub-MIC levels is possibly due to the total resistance assumption of our main framework. By the continuity of the effect function with respect to *zMIC*, the same trend is expected for sufficiently high partial resistance. To validate this expectation, we present numerical simulations of various partial resistance levels in the SI. Figures SI1, SI2, and SI3 also corroborate the superiority of the biostatic drug in suppressing resistance. Further, the figures show that low-level-resistance bacterial cells (i.e., *nMIC <* 2 *× zMIC*) are markedly suppressed at sub-MIC drug concentrations. In general, except for cases of high HGT rates, the survival probability is maximal at intermediate drug concentrations, consistent with previous results [1, 24]. In the next section, we consider dose-dependent mutations for in-depth insights into the mechanism of antibiotic-induced AMR.

**Figure 5.**
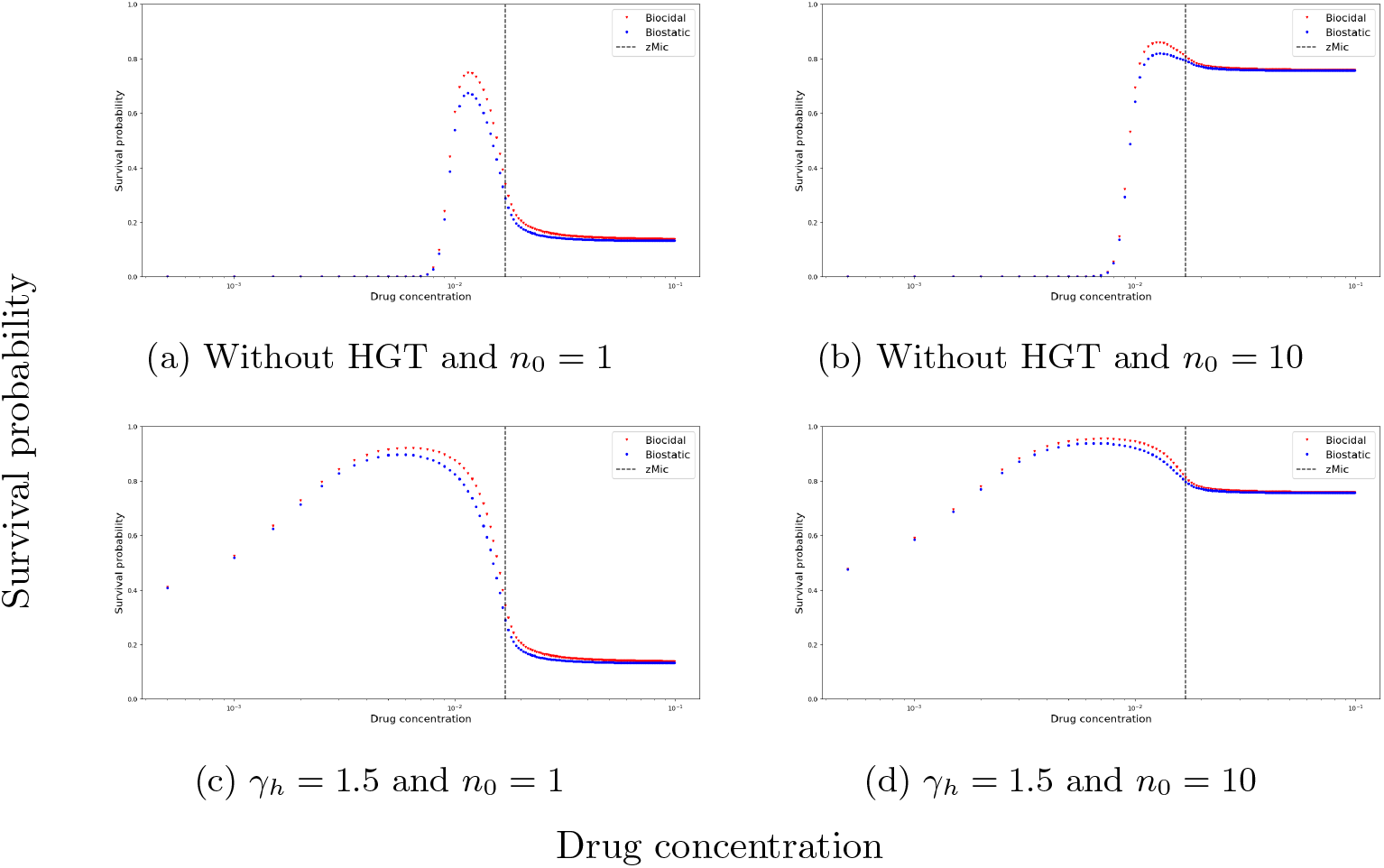
Survival probability under SGV with respect to drug concentration for 72-h PD biostatic vs. biocidal treatments with *K* = 10^7^ and *ϵ*_0_ = 10^−8^

### 4.2 Dose-Dependent Mutation

In this section, we analyze the effects of biostatic and biocidal drugs on various dose-dependent mutations for in-depth insights into the mechanism of antibiotic-induced AMR. As aforementioned, under deteriorating environmental conditions, bacterial organisms can benefit from a means to adapt, such as mutation [25]. This suggest a correlation between antibiotic treatment and bacterial population rescue via mutation. Although the literature on directed (non-random) mutation is somewhat controversial, recent studies have reported significant evidence of stress-induced mutation via various mechanisms, such as increasing the expression of error-prone DNA polymerases or down-regulating error-correcting enzymes [40, 41]. In terms of phenotypic mutation, it has been reported that *Pseudomonas aeruginosa*, a Gram-negative pathogenic bacteria that also exhibits notable levels of antibiotic resistance, changes shape in the presence of certain antibiotics that target cell-wall synthesis [42]. Recent experimental studies have reported evidence of the correlation of mutation rate and antibiotic concentration, particularly at sub-MIC levels [9, 43, 44]. Thus, we introduce a mathematical framework for antibiotic dose-dependent mutations. Recently, Teemu et al. [35] considered dose-dependent mutation and formulated a time-homogeneous optimal control problem. Meanwhile, besides our framework being time inhomogeneous, we approach this problem differently by investigating the interplay among competition release, HGT, SGV, and dose-dependent mutation.

#### 4.2.1 Linear Dose Dependence

Hongan Long et al. [43] reported a statistically significant increasing linear relationship between antibiotic concentration and mutation rate. In addition, Philip Ruelens et al. [9] reported a mutation that confers low resistance outcompeting a mutation that confers high resistance at low drug doses and vice versa, suggesting a decreasing trend with increasing drug concentration for the former mutation. Consequently, we explore three types of linear dose dependencies: (1) constant mutation rate, (2) linearly increasing rate, and (3) linearly decreasing rate. In general, we consider a linear function of drug concentration:

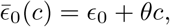

where *θ* = 0, *θ >* 0, and *θ <* 0 correspond to cases (1), (2), and (3), respectively. Notably, because we neglect backward mutation and hence 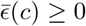 for all *c* ≥ 0, we set

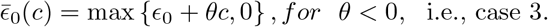

##### Corollary 1

*Suppose T <* ∞ . *Assuming E* ≡ *E*^∗^. *For any t* ∈ [0, *T*], *the following holds true*.

1. *If θ<0 and* ∈_0_ ≤ △^5^, *then* 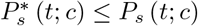.
2. *If θ<0 and* ∈_0_ ≥ △; *then 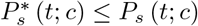, whenever* 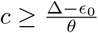.
3. *If θ<0 and* ∈_0_ ≤ △; *then* 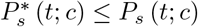, *whenever* 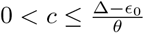.

#### 4.2.2 Nonlinear Dose Dependence

As is well-known that linear models mainly capture growth trends in data but fail to capture less-apparent features, we also consider the following nonlinear dose-dependent mutation rate:

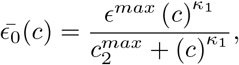

where *c* denotes the drug concentration, *ϵ*^*max*^ and 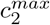 denote the maximum mutation rate and drug concentration to attain half of *ϵ*^*max*^, respectively, and *κ*_1_ determines the function’s slope. Although such nonlinear dose dependence has been considered for a polymyxin and peptide antibiotic [30], to the best of our knowledge, no such studies exist for ribosome-targeting antibiotics. The relevant parameters (i.e., *κ*_1_, *ϵ*^*max*^, and 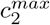) can be estimated from experimental studies for specific bacterial species following a previous study that employed an integrated approach to distinguish between collistin-induced and non-induced *Pseudomonas aeruginosa* resistance [30]. Albeit the absence of realistic parameters, we present a comprehensive theory and numerical simulations with hypothetical parameter values below. Moreover, we conduct a sensitivity analysis of these parameters with the ranges reported in the literature [30] (see SI).

##### Corollary 2

*Assuming E* ≡ *E*^∗^. *Suppose T <* ∞ . *For any t* ∈ [0, *T*], *the following holds true*.

1. *If ϵ*^*max*^ ≤ Δ, *then* 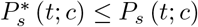;
2. *If ϵ*^*max*^ *>* Δ, *then* 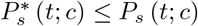, whenever 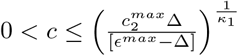 *where*

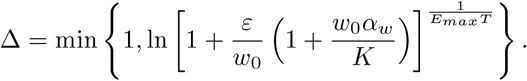

As shown in Figures 6 and 7, compared with the reference (constant mutation) cases, the more-realistic increasing linear mutation cases for our framework, in which total resistance is assumed, varies significantly for the biocidal treatment above *zMIC*. Figure 7 shows that a higher Hill coefficient leads to a lower sub-MIC peak of the survival probability. In addition, compared with the case of 1.1 Hill coefficient, that of coefficient 2 increases rapidly above the *zMIC*, indicating that aggressive biocidal treatment is detrimental to drugs with steep dose-dependent mutation rates. In general, increasing the drug concentration for biocidal treatment is ineffective in reducing the survival probability, suggesting that aggressively killing the sensitive bacterial population accelerates bacterial evolution toward resistance. Combined with Figure 3, these results indicate a trade-off in which faster clearance can be used to decrease the sensitive bacterial population size only at the expense of simultaneously increasing the chance of antibiotic-induced bacterial population rescue [35]. Under HGT and SGV, the qualitative trends of the dose-dependent mutation cases are similar to those of the reference cases (Figures 4 and 5).

**Figure 6.**
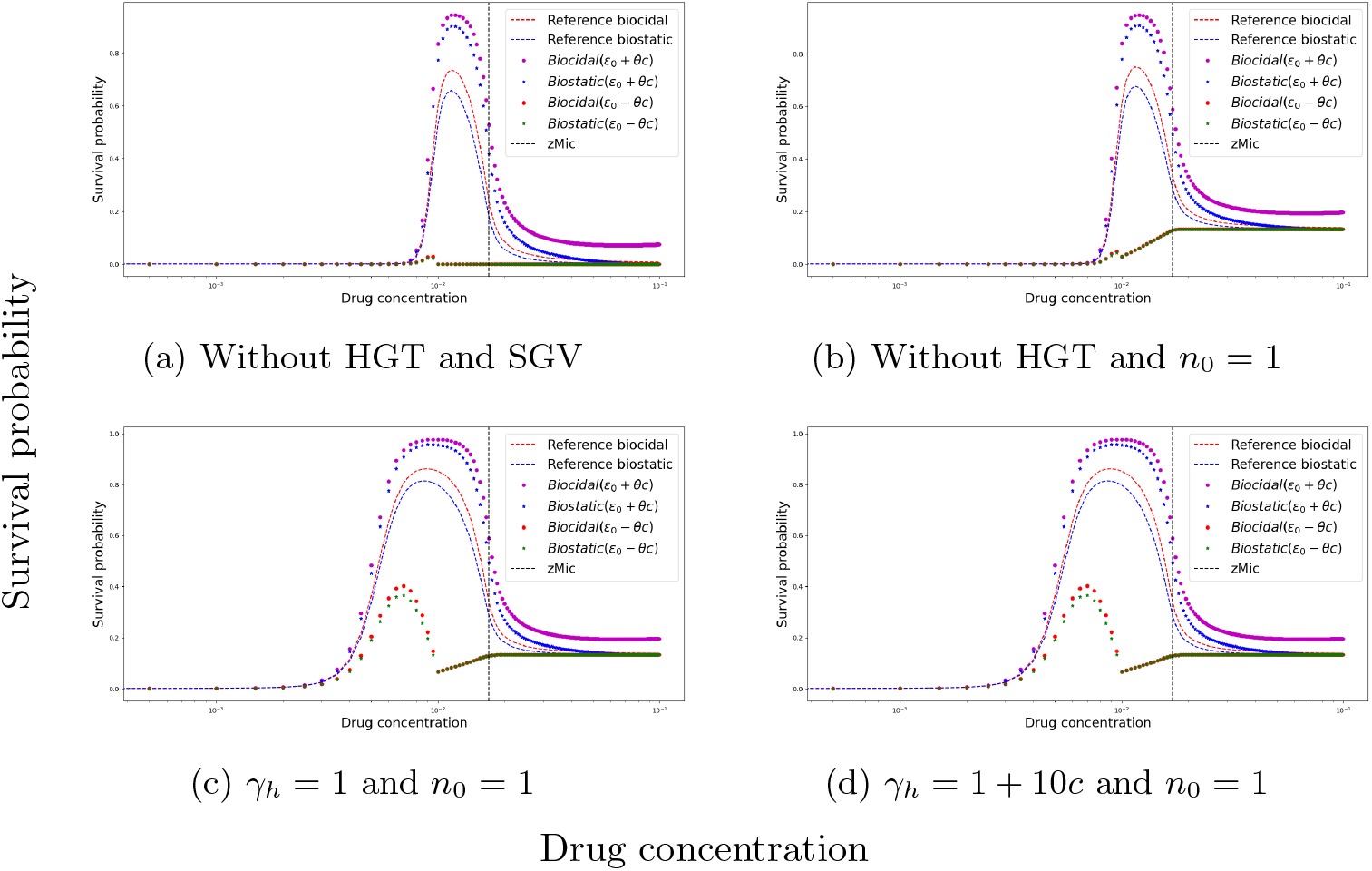
Comparison of various linear dose-dependent mutations for 72-h PD-based treatment with *K* = 10^7^, *ϵ*_0_ = 10^−8^, and *θ* = 10^−6^.

**Figure 7.**
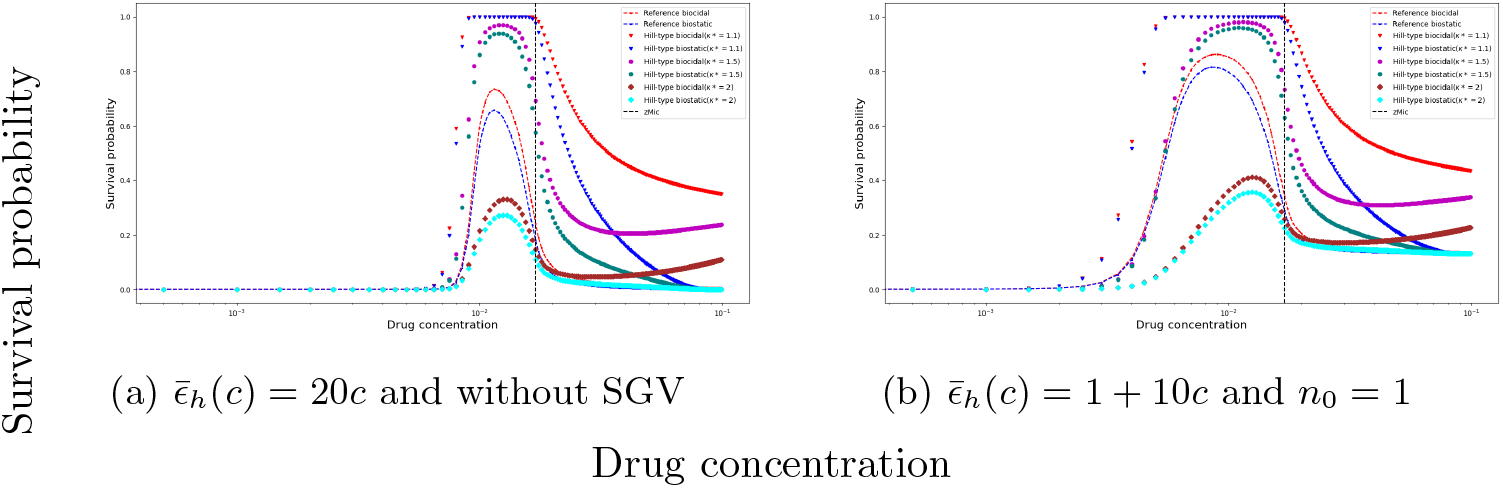
Survival probability with respect to drug concentration for 72-h PD biostatic vs. biocidal treatment considering various *de novo* dose-dependent mutation rates; *K* = 10^7^, 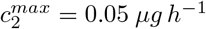, *ϵ*^*max*^ = 10^−6^, and *κ*_1_ = 1.1, 1.5, 2.

### 4.3 Effect of Nutrient Availability

As highlighted by Fernanda Pinheiro [29], antibiotic resistance is not only an evolutionary response problem but also an ecological response problem. Therefore, we analyze the effects of ecology, specifically nutrient availability, on bacterial susceptibility and resistance. Phenomenologically, nutrient-rich and nutrient-limited conditions in our model correspond to higher and lower carrying capacities, respectively. Consequently, we analyze the effects of carrying capacity on the wild-type strain’s susceptibility and the resistant strain’s survival.

#### 4.3.1 Finite Resources

We consider the carrying capacity as a variable parameter. Notably, unless specified otherwise, the following analyses are for both biocidal and biostatic.

##### Theorem 3

*Suppose K*_1_ ≤ *K*_2_. *For any t* ≥ 0, *the following holds true*.

- *w*^(∗)^(*t*; *c, K*_1_) ≤ *w*^(∗)^(*t*; *c, K*_2_)
- 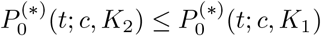
- 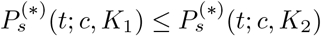

*Further, we have*

- 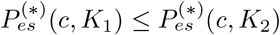

The above theorem indicates both higher antibiotic susceptibility and survival of resistant cells in a nutrient-rich environment than in a nutrient-limited environment. In addition, Theorem 3 suggests “a causality relationship between resource availability and antibiotic susceptibility and resistance.” We validate this theoretical result with numerical simulations (Figure 8). Perhaps, under nutrient-rich conditions, both sensitive and resistant bacteria tend to virulence. Competition between sensitive and resistant bacterial cells dominates the bacterial dynamics in a nutrient-limited environment, attributable to the well-known phenomenon of competitive release in which harsh treatment-induced selection pressure not only leads to the decay of sensitive cells but can also enhance the growth opportunities of the pre-existing or *de novo* resistant cells. This indicates that aggressive treatment accelerates bacterial evolution toward resistance because, by eradicating the competing sensitive cells, the resistant cells have even more resources to reoccupy the niche, leading to relapse [35], supporting the higher survival probability under biocidal treatment. Moreover, in the proof of Theorem 3 (Appendix F), we obtain that 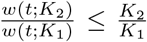 for wildtype populations not exceeding the carrying capacities. This inequality could inform how resource regulation affects the sensitive bacterial dynamics in the natural environment. We theoretically establish that reducing the number of sensitive bacterial cells via treatment while limiting available resources to avoid competitive release can prevent evolution toward resistance, consistent with the so called “ecological” intervention [45]. Further, transformative changes in the human diet or the use of probiotics can reduce nutrient availability to resistant bacteria while increasing antibiotic-sensitive bacteria.

**Figure 8.**
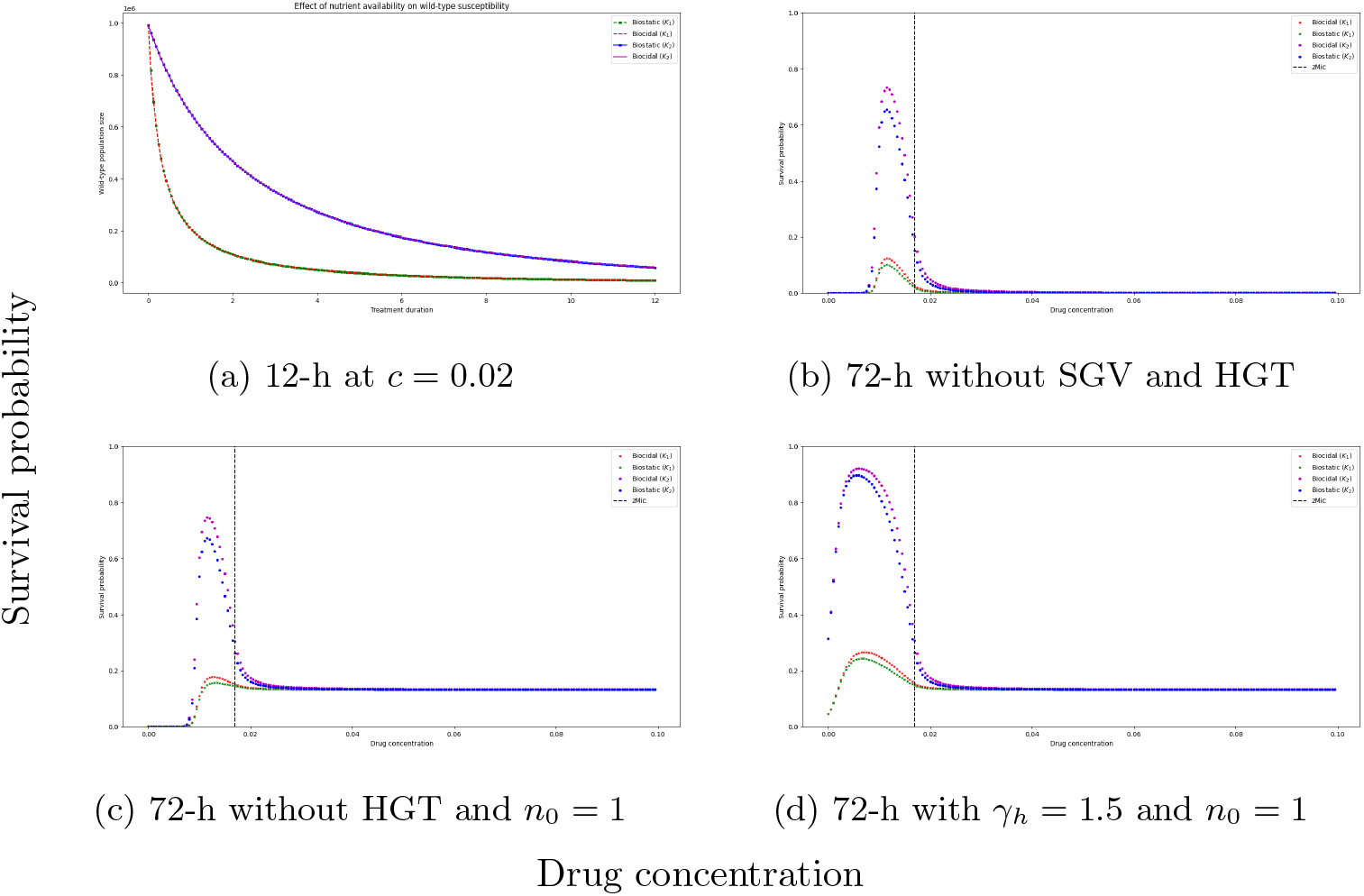
Effect of carrying capacity on (a) antibiotic susceptibility and (b–d) resistance survival with *ϵ*_0_ = 10^−8^; *K*_1_ = 10^6^ *< K*_2_ = 10^7^.

Below, we offer an optimal resistance-mitigation strategy via switching off birth with biostatic treatment in the absence of SGV. The following result (Theorem 4) indicates an “almost sure” extinction of *de novo* resistant strains with a “perfect” biostatic treatment (i.e., *E*^∗^ = *α*_*w*_).

##### Theorem 4

*Suppose ϵ*_0_ *>* 0. *If E*^∗^(*c*) = *α*_*w*_, *then* 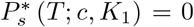 *for any K*_1_ *>* 0, *where the survival probability of de novo resistant strain* 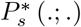 *is expressed as a function of the carrying capacity, K:*

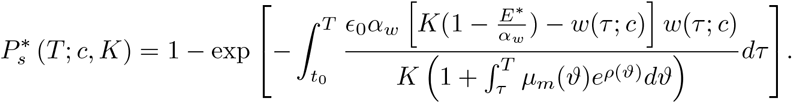

**Proof** Let *ϵ*_0_ *>* 0.

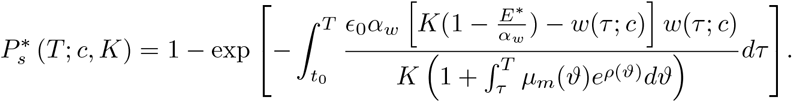

Setting *E*^∗^ = *α*_*w*_, we have

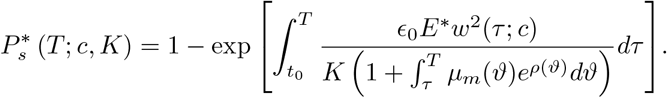

Thus,

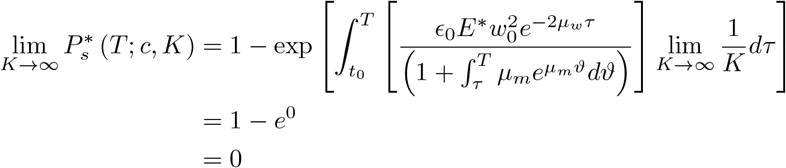

Because *K* → ∞, we can assume *K*_1_ ≤ *K*. Consequently, applying Theorem 3 and sandwiching, we obtain 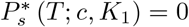.

#### 4.3.2 Infinite Resources

We phenomenologically associate an infinite resource environment with a bacterial population that grows exponentially [28]. Kindly refer to Appendix G for the analysis.

### 4.4 PK/PD: Incorporating Periodic Drug Intake

As in our model above, most existing related frameworks [1, 7, 8, 10] assume constant drug concentration, an assumption that is unrealistic for non-intravenous treatments. Such an assumption is often made for one or both of the following reasons: first, the effect of variation in drug concentration is often sufficiently small that it can be neglected; and, second, models with constant drug concentration are more tractable. However, if the effect of variation in drug concentration is not sufficiently small, any predictions based on a model that fails to account for this variation could be highly inaccurate. In this situation, tractability needs to be improved. Following G. A. Koch-Noble [46] and Andrea McNally [47], we consider a one-compartment ODE model for periodic antibiotic intake:

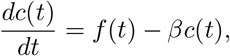

where *c*(*t*) is the drug concentration at time t, *β* is the drug metabolism rate, and *f* (*t*) is the dosing pattern of drug intake (a discrete function, defined using the Dirac delta function):

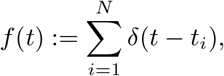

where *N* is the number of times the drug has been taken. At the end of the treatment course, the solution to the above ODE is given by [47]

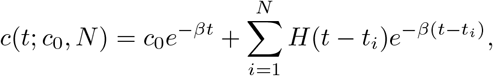

where *c*_0_ is the initial (maximum) drug concentration and 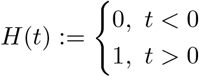 is the distributional antiderivative of *δ*(*t*). We now incorporate the PK model into the Regoes’ PD model [32] to obtain the following PK/PD-based model:

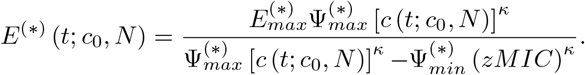

Similarly to the PD-based cases, except for the time dependency of the PD effect function, we solve for the expected wild-type dynamics (summarized in Table 4).

**Table 4.**
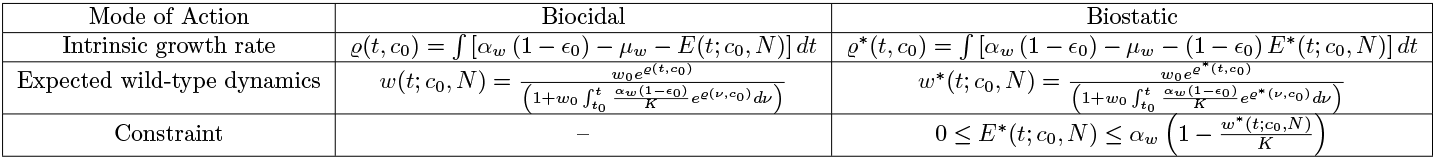
Wild-type dynamics for PK/PD-based biocidal vs. biostatic.

**Table 5.** Wild-type dynamics of single PK/PD-based biocidal vs. biostatic treatment dose.

| Mode of Action | Biocidal | Biostatic |
| --- | --- | --- |
| | $\varrho_i(t, c_0) = \int [\alpha_w (1 - \epsilon_0) - \mu_w - E_i(t, c_0)] dt$ | $\varrho_i^*(t, c_0) = \int [\alpha_w (1 - \epsilon_0) - \mu_w - (1 - \epsilon_0) E_i^*(t, c_0)] dt$ |
| | $w_i(t; c_0, w_{i,0}) = \frac{w_{i,0} e^{\varrho_i(t, c_0)}}{(1 + w_{i,0} \int_{t_i}^t \frac{\alpha_w (1 - \epsilon_0)}{K} e^{\varrho_i(\nu, c_0)} d\nu)}$ | $w_{i,0}^*(t; c_0, w_{i,0}^*) = \frac{w_{i,0}^* e^{\varrho_i^*(t, c_0)}}{(1 + w_{i,0}^* \int_{t_i}^t \frac{\alpha_w (1 - \epsilon_0)}{K} e^{\varrho_i^*(\nu, c_0)} d\nu)}$ |
| | – | $0 \leq E_i^*(t; c_0) \leq \alpha_w \left(1 - \frac{w_i^*(t; c_0, w_{i,0}^*)}{K}\right)$ |

We further assume a uniform period of treatment, i.e., *t*_*i*+1_ − *t*_*i*_ = Π and require that *c*_0_ = *c*(*t*_1_ = 0) = *c*(*t*_2_ = *t*_1_ + Π) = …. = *c*(*t*_*N*_ = *t*_1_ + (*N* − 1) Π). Although this assumption is rationalized for drugs with fast metabolism rates (half life substantially less than the dosing interval) [48] and those administered at low doses, it may, at first, seem irrational for drugs administered at high doses or drugs with slow metabolism rates. However, considering that some ribosome-targeting antibiotics, e.g., ciprofloxacin, can be excreted via feces, urine, or sweat [49], the above assumption is rational. In particular,

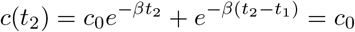

Yields

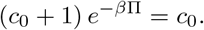

Solving for Π, we obtain

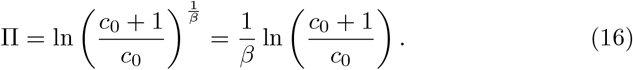

For 1 ≤ *i* ≤ *N* and *t* ∈ [*t*_*i*_, *t*_*i*+1_), we consider

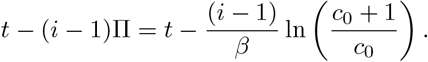

Hence,

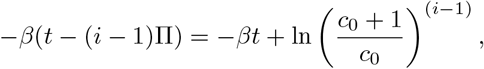

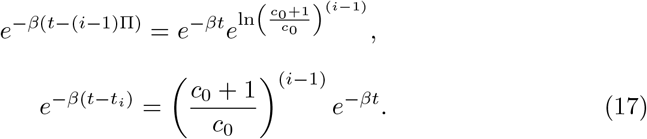

Therefore, for the *n*-th treatment course, we have

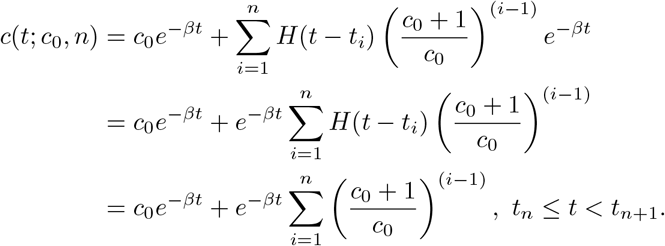

Using the geometric sum formula, we obtain the following:

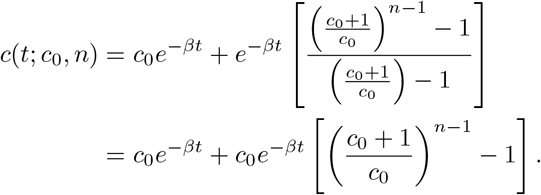

Thus, for the *n*-th treatment course, we obtain the following explicit PK model:

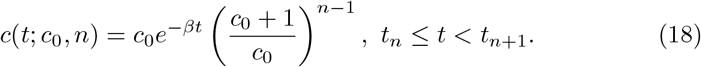

Substituting Eq. 16, we have

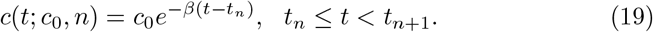

Before forging ahead, it is worth highlighting that, realistically, the PK dynamics (Eq. 19) and proposed GBDP-I operate on different time scales, with the former being slower. Biologically, this is motivated by the fact that synaptic plasticity is slower than neural activation [50], which can, respectively, be associated with metabolism and cell growth. However, because the PK model has only one independent rate parameter, *β*, we can handle the time-scale discrepancy in the PK/PD-based model by re-scaling *β* to be relatively small such that the timescale factor can be set to unitary without ambiguity. Notably, Eq. 19 is similar to the PK equation proposed by Regoes et al. [32], justifying its compatibility with the classical PD model (Eq. 3). Moreover, *β* can be estimated based on the classical experiment of administering drug, measuring plasma concentration over time, and fitting scatter plots, from which the adequate dosing schedule Π can be inferred (see, e.g., [51] and references therein). In addition, owing to the periodicity of the PK model, it is sufficient to analyze the dynamics of the PK/PD model in one course of treatment, say, within [*t*_*i*_, *t*_*i*+1_), denoted as *E*_*i*_:

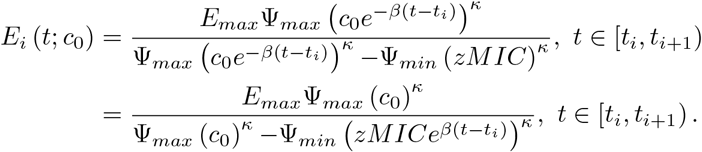

We now consider the dynamics of *w*(*t*; *c*_0_) within each treatment course, [*t*_*i*_, *t*_*i*+1_), denoted as *w*_*i*_(*t*; *c*_0_, *w*_*i*,0_), with *w*_1_(*t*_1_; *c*_0_) := *w*_0_ ≜ *w*_1,0_ and 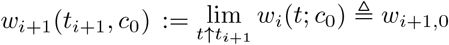, for *i* = 1, 2, · · ·, *N*.

Further, to enhance simulation tractability, we manually compute the intrinsic growth rates. For instance, for the biocidal drug effect, we have

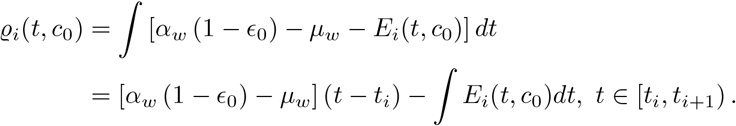

Moreover,

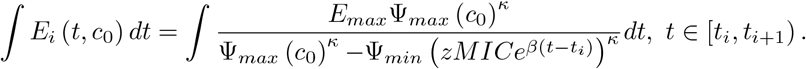

Let *A* = *E*_*max*_Ψ_*max*_ (*c*_0_)^*κ*^, *B* = Ψ_*max*_ (*c*_0_), *C* = −Ψ_*min*_ (*zMIC*), and *ν* = *βκ*.

Thus,

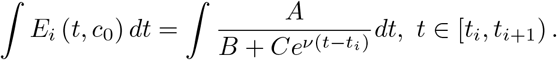

Integrating by substitution and partial fraction, we obtain

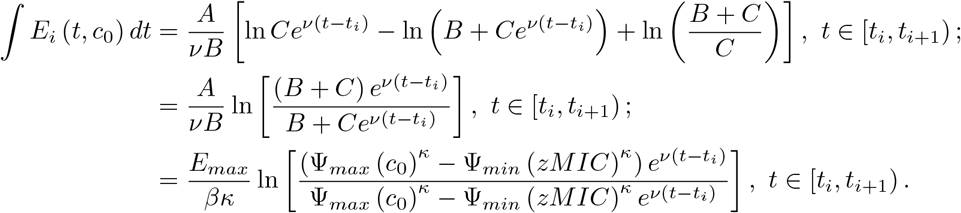

The following theorems establish analytical results comparable to the above PD-based analytical results with PK incorporation.

#### Theorem 5

*Suppose E* ≡ *E*^∗^. *Then, w*(*t*; *c*_0_, *N*) ≤ *w*^∗^(*t*; *c*_0_, *N*), ∀*t* ≥ 0, *N* ∈ ℕ.

**Proof** Let *N* ∈ ℕ and *ϵ*_0_ *>* 0 be fixed. Assume *E* ≡ *E*^∗^. Thus, 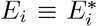 for each *i* = 1, 2, *…, N* . Thus, we have that 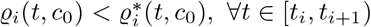. In particular, 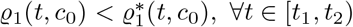.

Therefore, by the basic comparison theorem of ODEs, we have

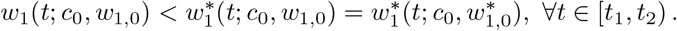

Taking limit as *t* → *t*_2_, we have that 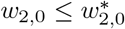. Moreover, because 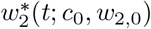 and 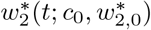 have the same growth rate, 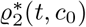, we obtain

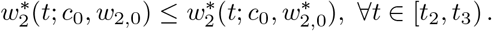

Again, by the basic comparison theorem of ODEs

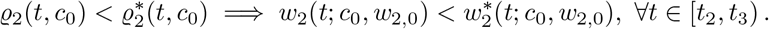

The above two inequalities yield

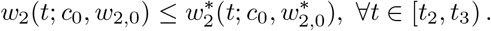

It then follows by induction that, for each *i*,

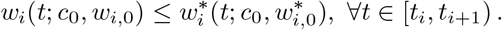

Thus,

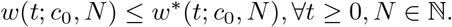

#### Theorem 6

*Assuming E* ≡ *E*^∗^. *Let t*_*N*_ = *T <* ∞. *Then*, ∃ Δ *>* 0 *such that* 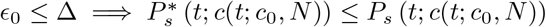, ∀*t* ≥ 0, *where*

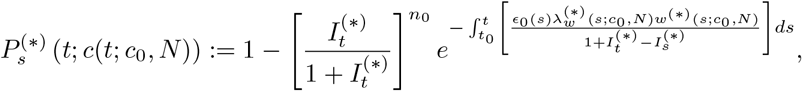

*Moreover, we can estimate* Δ *as follows:*

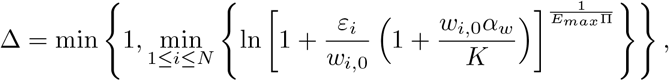

*where* Π *denotes the treatment period and ε*_*i*_ *is a chosen bound on the maximal wild-type density difference for the i-th treatment course*.

**Proof** Because *N* is finite and *w*^(∗)^(*s*; *c*_0_, *N*) is piecewise continuous, we merely apply Theorem 2 for each piece [*t*_*i*_, *t*_*i*+1_), *i* = 1, 2, · · · *N*.

#### 4.4.1 PK/PD-Based Simulation

For simulations to assess the effect of PK on the resistant bacterial evolution, in general, we can compare a baseline treatment strategy (PD case) with PK/PD treatment strategies in terms of the area under the concentration–time curve (AUCTC). First, we fix a final treatment duration of *T* . Second, for a fixed initial drug concentration, *c*_0_, we consider a Π-periodic dosing strategy prescribed to be administered in 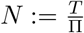 courses and determine the corresponding constant concentration, *c*, such that the AUCTC of each treatment period matches the baseline PD’s AUCTC (i.e., 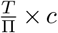). Consequently, the baseline PD’s AUCTC is *T × c*. Moreover, the corresponding metabolism rate for each periodic treatment strategy is given by Eq. 16. Mathematically, we obtain the following:

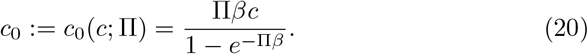

Further, the combination of Eqs. 16 and 20 yields

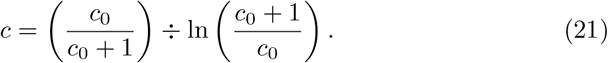

From Figure 9, albeit the consistency in qualitative trends with the baseline PD-based model, it is evident that the PK dynamics considerably influences the bacterial evolution. Particularly, PK incorporation increases the difference between the biostatic- and biocidal-based survival probabilities of the resistant strain, validating the efficacy of a biostatic drug in resistance suppression. The 12-h dosing case yields higher survival probabilities than constant efficacy within the 72-h treatment window for both types of drugs and initial mutant population sizes. This result suggests that a faster decay in drug concentration makes it so that the drug efficacy more quickly becomes below the amount to suppress wild-type reproduction, and this is repeated every period, so that the wild-type population remains, allowing for mutations to resistant strains, which can also survive.

**Figure 9.**
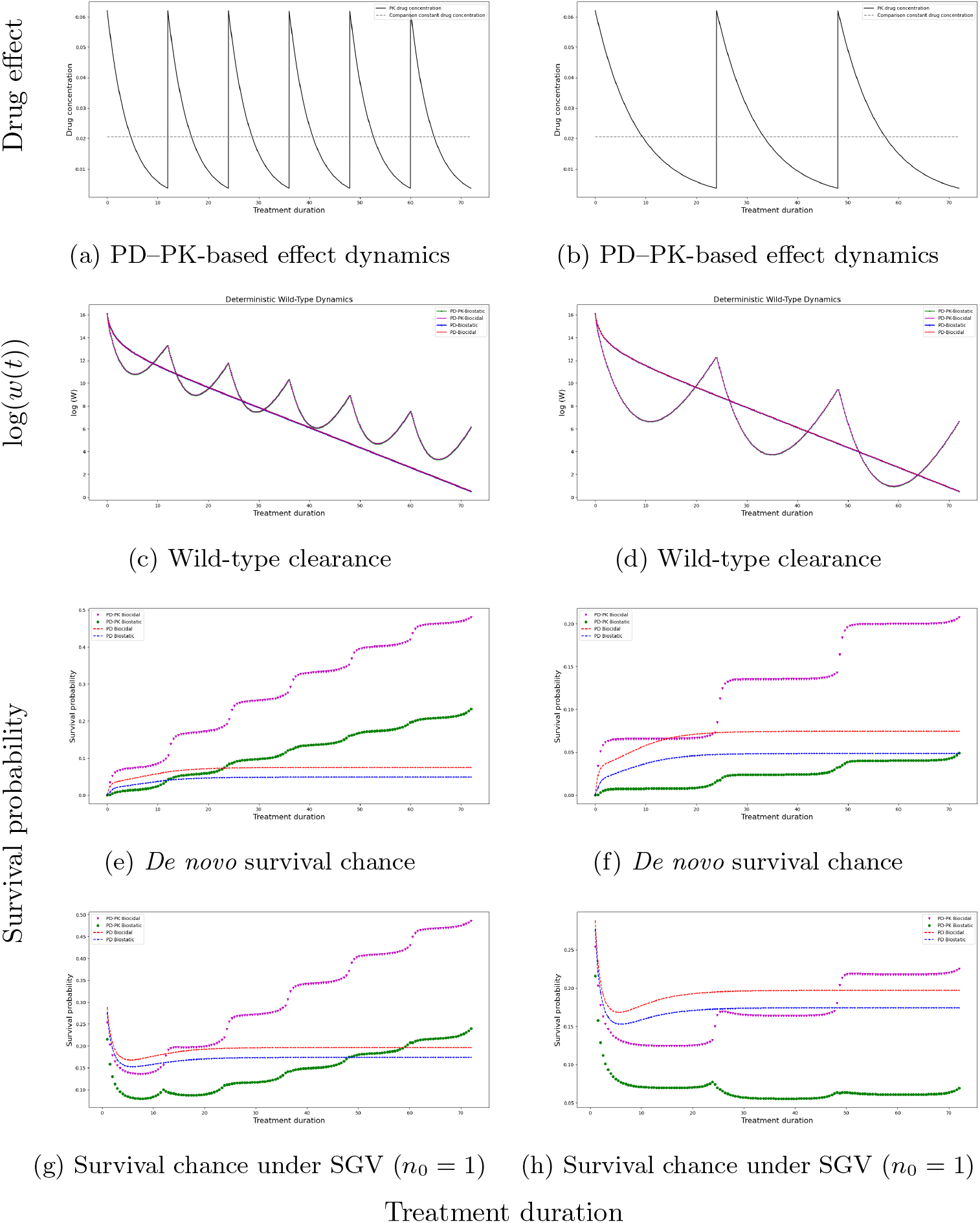
Comparison of 12-h and 24-h periodic dosing with matched AUCTC constant concentration.

Interestingly, within the considered 72-h treatment duration, the biostatic 24-h periodic case, especially under SGV, shows a lower survival probability than both the 12-h dosing and PD-based cases. In particular, the biostatic 24-h periodic dosing under SGV exhibits a decreasing survival probability of resistant cells over the majority of the 72-h treatment duration (Figure 9h). Thus, administering a biostatic drug can be less sensitive to variations in concentration, even at sub-MIC levels for relatively long time periods, with respect to suppressing resistance establishment. Taken together, these results suggest that besides the drug mode of action and concentration, the time dependency of drugs should be considered for effective use against AMR.

To further explore how temporal variations in drug concentration affect resistance establishment, we present simulations of PD-based and PK/PD-based survival probabilities over extended treatment durations in Section SI 4. Figure SI 9 demonstrates that the baseline PD-based survival probabilities saturate at short duration, unlike the PK/PD cases, indicating the need for an in-depth understanding of the effect of PK dynamics on AMR. Particularly, the PD-based survival probability converges for *T <* 72 *h*. Meanwhile, there is an average increasing PK/PD-based survival probability for the considered longer treatment duration (*T* = 300 *h*), indicating convergence over a longer time horizon, as probability values are bounded above by one. The larger values of survival probability obtained reflect the periodically rebounding wild-type population during dips in drug concentration that increase the probability of resistance mutations. However, a rationale for the validity of our simplified model is that the wild-type bacteria population can be considered deterministic, which may be violated under the periodic fluctuations, particularly for the 24-h dosing scheme, where the wild-type population reaches very low densities that might cause extinction before the population can rebound when drug efficacy dips. In addition, PK incorporation widens the difference between the biocidal and biostatic survival probabilities, suggesting that patient-specific PK measurements obtained through therapeutic drug monitoring are critical for accurately predicting resistance emergence and guiding personalized antibiotic dosing strategies.

## 5 Discussion

We study within-host bacterial dynamics using a CTMC. Particularly, we propose a PD-based CTMC model considering HGT, SGV, and the *de novo* emergence of a resistant bacterial strain via spontaneous or drug-induced mutation as well as analyze the effects of drug mode of action and nutrient availability on antibiotic resistance and susceptibility. Making some biological plausible assumptions, we approximate the resistant sub-CTMC with a GBDP-I for tractability. Notably, the GBDP-I approximation is based on a corresponding nonlinear ODE, unlike other techniques that approximate with the linearized version of the (nonlinear) system of ODEs at equilibria [52]. Based on the GBDP-I, we analyze the survival chance of the resistant bacterial strain under biostatic and biocidal treatment conditions. Moreover, we analyze the deterministic clearance rate of the sensitive bacterial strain, as the sensitive sub-CTMC is well-described by its deterministic ensemble [1]. In addition, we rigorously analyze the effects of nutrient availability on antibiotic susceptibility and resistance.

The findings of this study suggest that biocidal treatment is more effective in eradicating sensitive bacterial strains whereas biostatic treatment is superior in suppressing resistance. Although our main analysis is based on full resistance (i.e., *rMIC* = ∞ [1]), simulations considering partial resistance^6^ show the same qualitative trend (see SI). Moreover, incorporating PK is essential because survival probabilities differ markedly from the constant case and often increase substantially with treatment duration, especially under biocidal treatment. Overall, simulations considering PK or low partial resistance levels indicate markedly lower survival probabilities of the *de novo* resistant strain with biostatic treatment at relatively low drug concentrations. Thus, we infer that a biocidal drug is more suited for the “hit hard, hit early” strategy whereas administering a biostatic drug at low doses suppresses resistance significantly, consistent with previous experimental and numerical studies (see, e.g., [1, 39]). Besides debunking the “one size (style) fits all” notion, the analytical findings, as well as numerical validations, of this study highlight the need for treatments that combine drugs with different drug modes of action for synergy, facilitating personalized treatment strategies via therapeutic drug monitoring [53] and combination therapy [10]. Notably, the primary goal of our numerical simulations is not to provide quantitative predictions for a specific disease, but rather to validate and explore the qualitative behavior of the complex AMR dynamics.

This study challenges the conventional inquiry of the superiority between the two extremal dosing strategies to mitigate AMR and instead analyzes the ideal extremal dose for a specific drug mode of action. Although the deterministic bacterial dynamics are the same for the two modes of action when the antibiotic concentration is sufficiently small [1], our stochastic analyses indicate a significant difference in AMR suppression. The simple and robust theoretical result of the superior susceptible bacterial clearance effect of biocidal drugs at high drug concentrations is not even verified in clinical settings. Some meta-analyses found no difference in treatment success between biostatic and biocidal drugs [37, 38]. It is possible that drugs that are theoretically biostatic, in practice, also directly kill bacteria at clinically relevant doses [39]. Nevertheless, Figures 4–7 show that a high supra-MIC biocidal concentration is required for resistance suppression. This suggests that biocidal drugs should be administered at sufficiently high doses and more frequently to ensure bacterial clearance before resistance-induced population rescue occurs. Notably, biostatic treatment is a more suitable choice for bacterial infections with high mutation rates. Further, for enhanced treatment outcome, combination therapy may be better to synergize the effects of both drug modes of action. Particularly, we hypothesize that a high-biocidal–low-biostatic combination therapy is more suitable for sensitive bacterial eradication without resistance-induced rescue than a low-biocidal–high-biostatic combination therapy. A theoretical study based on this hypothesis is ongoing. As a promising next step, *in vitro* experiments should be conducted to test the impacts of different modes of action on AMR evolution and characterize the probability of emergence of AMR depending on drug concentration and drug mode of action. The pronounced advantage of a biostatic drug in suppressing *de novo* resistance suggests a “greener” treatment than its biocidal analog, as this suppression prevents environmental contamination or secondary infection by resistant bacteria.

Recently, Fernanda Pinheiro [29] posed the following question “if the abundance of the antibiotic target depends on the growth condition, does the same happen to antibiotic susceptibility?” Although the proposed model is not fully mechanistic, we answer this question in the affirmative by associating nutrient-rich and nutrient-limited environments, respectively, with high and low carrying capacities of logistic growth—a standard practice in mathematical biology. The results suggest that combating resistant bacteria, as well as sensitive bacteria, with probiotics to reduce accessible nutrients and hence increase competition can mitigate antibiotic resistance, highlighting the critical role of ecological competition in shaping the evolution of antibiotic resistance. In this context, resistance evolution may be mitigated when resistant bacteria are unable to exploit ecological niches vacated by sensitive populations. From a clinical perspective, our results on the effect of nutrient availability further suggest that interventions targeting the microbial environment may complement conventional antimicrobial therapies. Particularly, substantial dietary modifications or the use of probiotics could alter nutrient dynamics within the host microbiome, thereby reducing the resources accessible to resistant bacteria while killing the sensitive sub-population. Although the use of probiotics cannot eradicate pathogenic bacteria, it can increase competition of the declining sensitive bacterial subpopulation and *de novo* or pre-existing resistant bacterial cells, thereby delaying evolution toward resistance. Such ecological approaches may contribute to preserving microbial competition and preventing resistant strains from becoming dominant. Future transdisciplinary research should exploit combining resource-limiting strategies [45] with traditional ribosome-targeting antibiotics, as well as transformative changes in human diet for enhanced immune response, to combat antibiotic resistance.

Coates et al.[54] reported that biocidal treatment clears the bacterial population size faster than biostatic treatment. Moreover, Alexander and MacLean [55] reported that higher SGV increases the probability of survival of the resistant strain. Although some of our findings have, in fact, already been tested experimentally, our theoretical work motivates several interesting experiments, such as studying how nutrient availability shape the fitness landscape of multi-strain bacteria as well as how drug resistance evolves from no/small population(s) of resistant cells during treatment, especially for drug-inducible resistance, even without concentrations that give the resistant subpopulation a fitness advantage. Particularly, *in vitro* time–kill experiments could be conducted to characterize the probability of emergence of *de novo* resistance depending on drug concentration, nutrient availability, and the drug mode of action. We predict that the differences between the two drug modes of action are more pronounced under a nutrient-rich environment and when PK is incorporated. The validity of our prediction on the drug-induced *de novo* resistant survival probability can also be assessed. To this end, besides estimating HGT rates from conjugation assays, one needs to expose a bacterial population comprising only sensitive cells to antibiotics and empirically determine the rate of phenotypic/genotypic changes, if any, as well as measure the survival probability of phenotypic/genotypic-mutated cells as a function of the antibiotic concentration.

In parallel, one can evaluate all terms of the dose-dependent mutation functions through simple *in vitro* time–kill experiments and scatter-plot-fitting. The dose-dependent mutation parameters can all be measured *in vitro* at different antibiotic concentrations following procedures outlined in [30]. Our prediction that biostatic drugs are better than biocidal drugs in suppressing resistance evolution when the mutation rate is below a certain threshold can even be tested without knowledge of the dose-dependent mutation parameters, provided one accounts for the stochastic nature of resistance evolution. Stochasticity implies that not all phenotypic/genotypic mutations will found a lineage that survives or becomes established. Although these experiments would help validate our theoretical predictions, their implementation and systematic evaluation are beyond the scope of this study and are left for future work.

In conclusion, we propose a PK/PD-based CTMC model that explicitly captures *de novo* bacterial strain emergence. Using this framework, we investigate how distinct drug modes of action, mutation mechanisms, nutrient availability, and drug concentration variability shape antibiotic resistance evolution. By approximating a multi-component stochastic model with a birth–death– immigration process, we provide an analytical framework for studying resistance emergence and generating experimentally testable predictions. Future work will focus on validating the predictions through comparison with experimental measurements of resistance emergence under antibiotic treatment.

## Data Availability

All code used in this study is publicly available at https://github.com/chimezie0124/denovo_amr_prob_for_biostatic-biocidal.

## Appendix A Notations

**Table 6.**
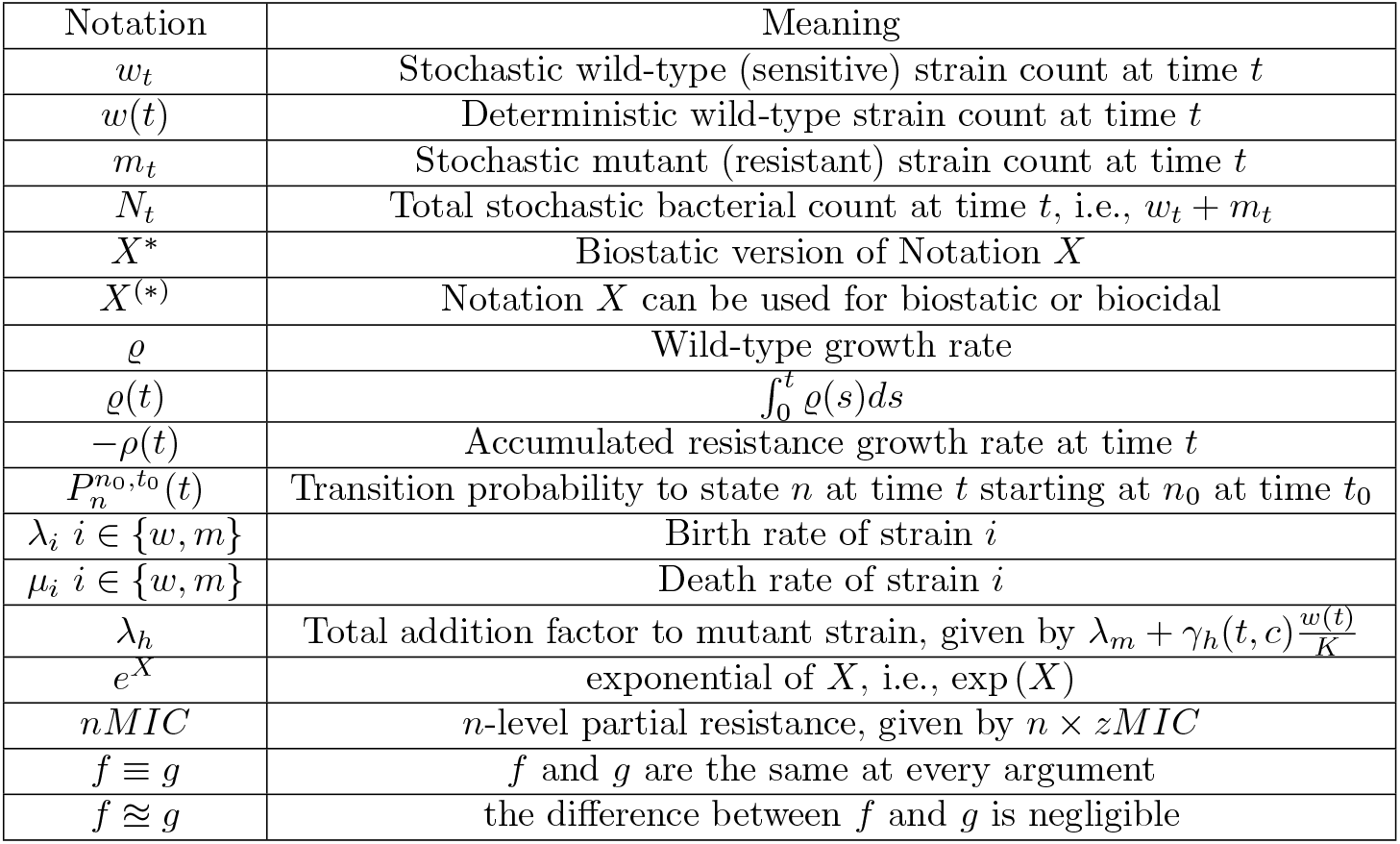
Table of notations.

## Appendix B PDE Solution

Based on the method of characteristics, we set 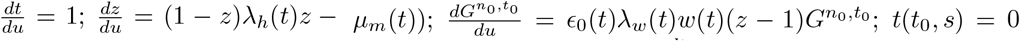; *z*(*t, s*) = *s*; and 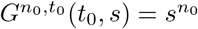. Because 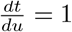, we obtain the following ordinary differential equations (ODEs):

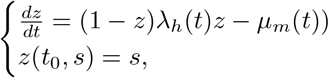

and

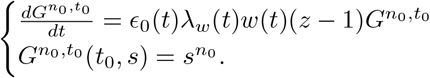

We now consider

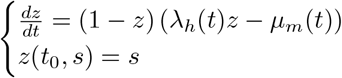

and make the substitution 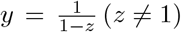, which yields the following linear initial value problem:

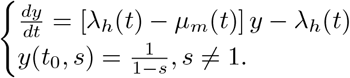

Employing the integrating factor, 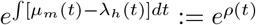, we obtain

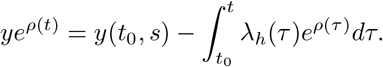

Recovering the initial substitution, we obtain

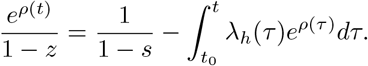

Solving for *s*, we obtain

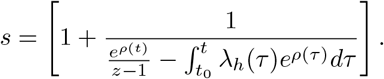

Therefore, solving the second initial value problem, we obtain

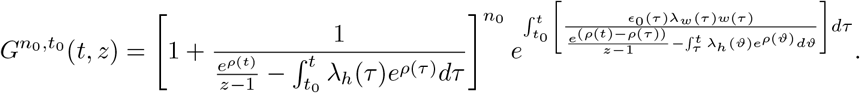

## Appendix C Proof of Proposition 2

**Proof** Define

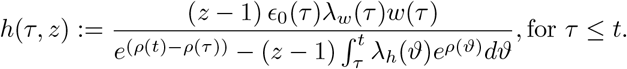

Expanding as a power series of *z*, we obtain

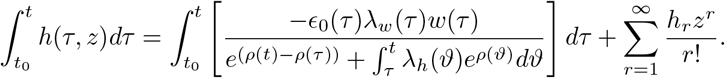

Thus,

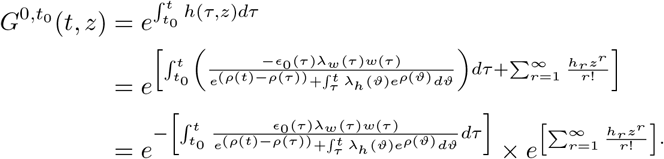

Because 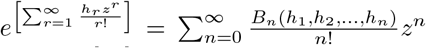, where *B*_*n*_’s are the complete

Bell polynomials [27], we obtain

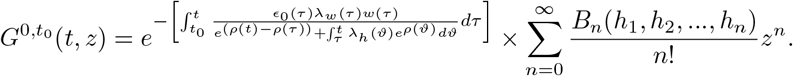

From the terms of this series, we obtain

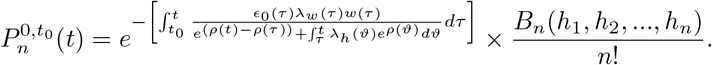

Thus, setting *n* = 0, we obtain the extinction probability of *de novo* mutations as follows:

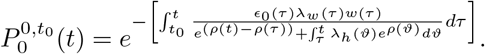

## Appendix D Proof of Theorem 2

**Proof** Because *E* ≡ *E*^∗^ and *T <*, uniform continuity of the given ODE’s solution with respect to the parameters yields the existence of Δ *>* 0 such that *ϵ*_0_ ≤ Δ implies *w*(*t*; *c*) ≊ *w*^∗^(*t*; *c*) for any *t* ∈ [0, *T*]. Consequently, *ρ*(*t*) ≊ *ρ*^∗^(*t*) because

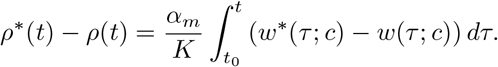

In addition, 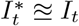 because

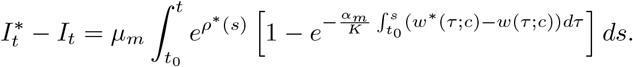

Therefore, for any *t* ∈ [0, *T*],

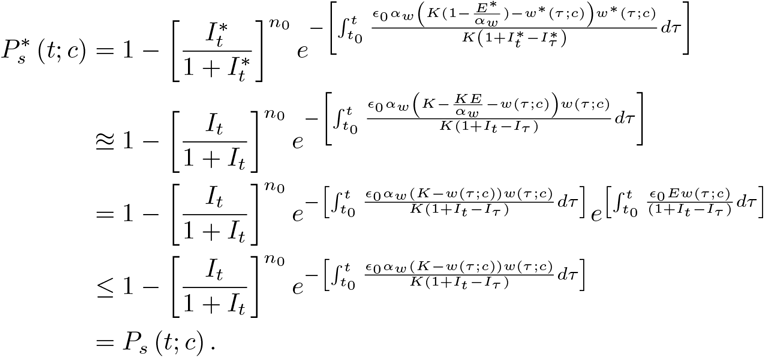

We now estimate Δ. Clearly, we must have that 0 *<* Δ ≤ 1; else, it is not a feasible bound for *ϵ*_0_.

For each *t* ∈ [0, *T*] and *ε >* 0, we seek Δ_*t*_ := Δ(*t, ε*) *>* 0 such that

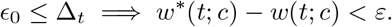

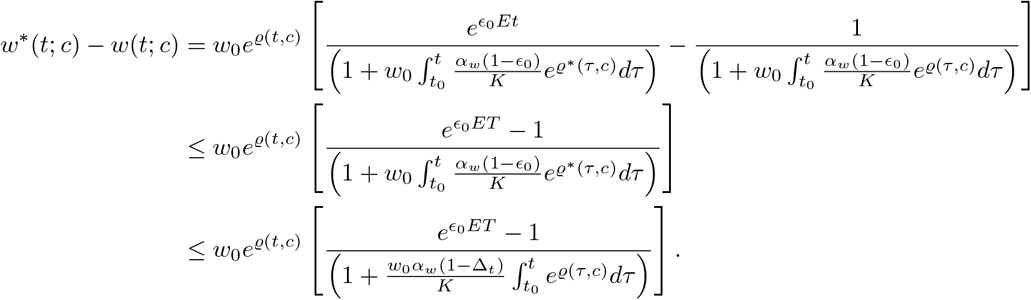

In addition,

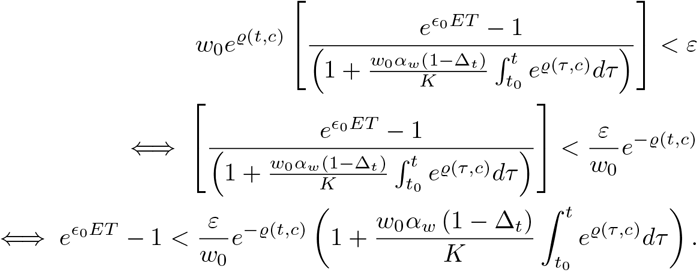

Further, 0 *< ϵ*_0_ *<* Δ_*t*_ ≤ 1 =⇒ 1 − Δ_*t*_ *<* 1. Consequently, we have

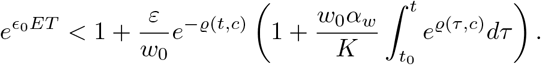

Thus,

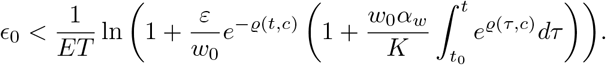

Therefore, we define

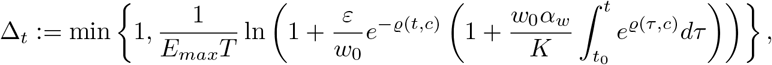

We now define Δ as follows:

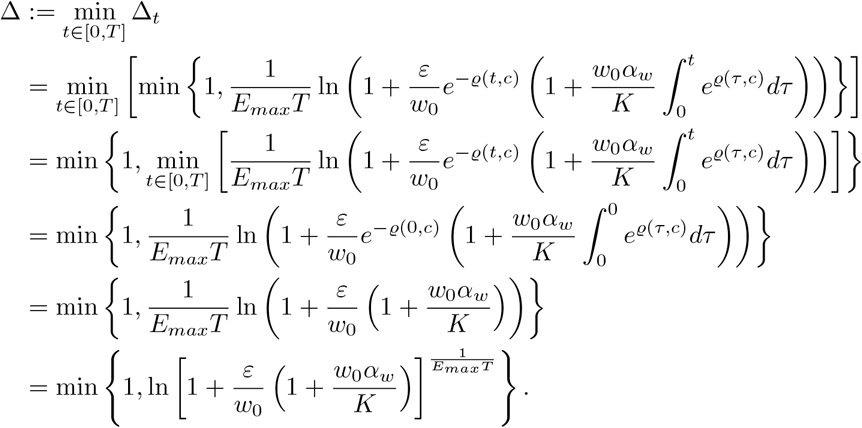

## Appendix E Proof of Corollaries 1 & 2

### E.1 Corollary 1

**Proof** It suffices to show that 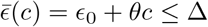, where

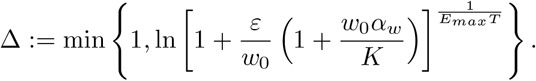

1. Let *θ <* 0. If *ϵ*_0_ ≤ Δ, then

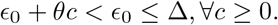
2. Again, let *θ <* 0. If *ϵ*_0_ ≥ Δ, then

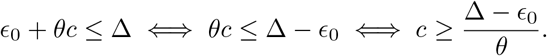
3. Finally, let *θ >* 0. If *ϵ*_0_ ≤ Δ, then

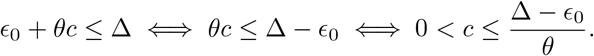

### E.2 Corollary 2

**Proof** We need only show that the assumptions of Theorem 2 are satisfied.

1. The proof follows directly from Theorem 2.
2. Notably, we have the following estimates:

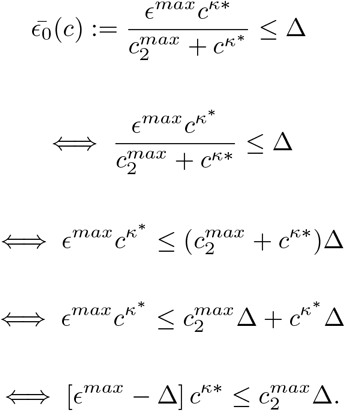

Thus, if *ϵ*^*max*^ *>* Δ, we have

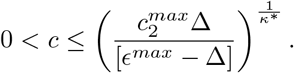

[*ϵ*^*max*^ − Δ] It then follows from Theorem 2.

## Appendix F Proof of Theorem 3

**Proof** *w*^(∗)^(*t*; *c, K*_1_) ≤ *w*^(∗)^(*t*; *c, K*_2_) follows directly from

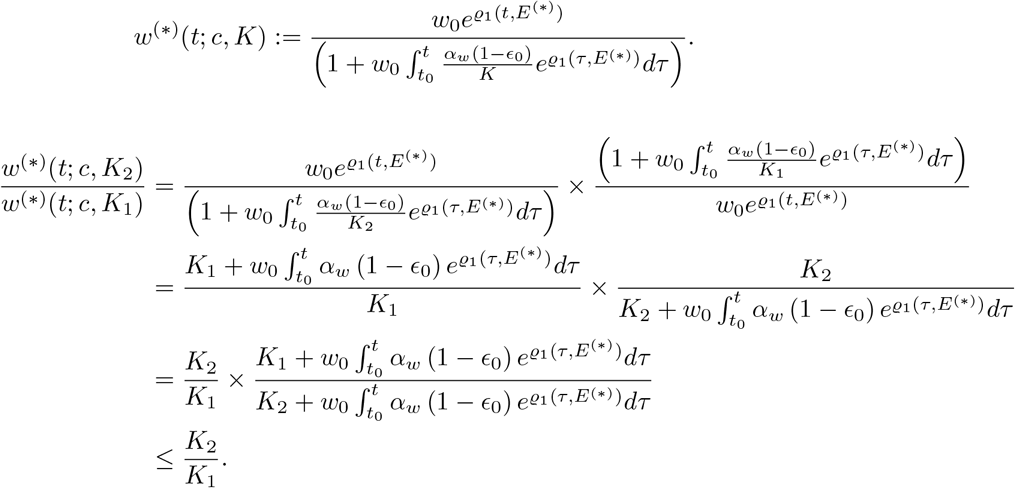

Thus,

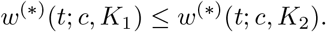

Further,

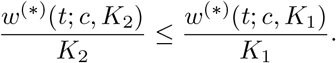

Therefore,

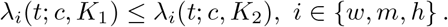

Consequently,

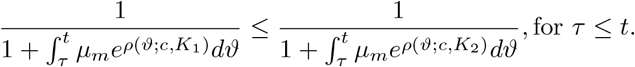

Thus,

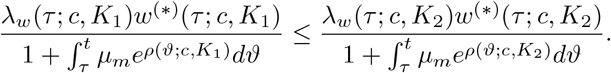

Therefore,

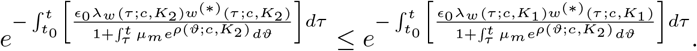

In addition,

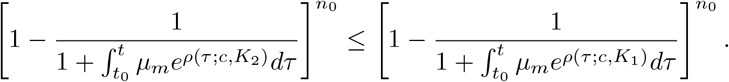

Hence,

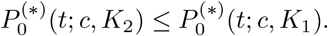

In addition, we obtain

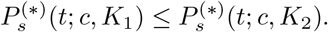

Further, taking limit as *t* → ∞, we obtain

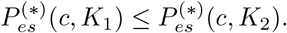

## Appendix G Infinite Resources

In this section, we consider a bacterial population that grows exponentially, which is phenomenologically associated with infinite resources [28]. That is, *λ*_*i*_(*t, N*_*t*_, 0) = *α*_*i*_ and *µ*_*i*_(*t, N*_*t*_, 0) = *µ*_*i*_, *i* ∈ {*m, w*}. Thus, *λ*_*h*_(*t, N*_*t*_, *c*) = *α*_*m*_ + *γ*_*h*_(*t, c*)*w*_*t*_.

### Biocidal

For biocidal treatment, the drug-mediated growth rate is given by

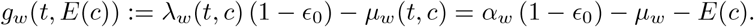

Consequently, we have

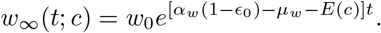

### Biostatic

For biostatic treatment, the drug-mediated growth rate is given by

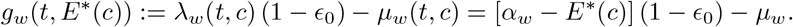

However, for biological relevance, we require that

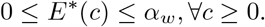

Under this constraint, we have

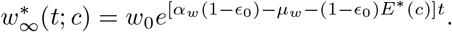

#### Theorem 7

*Suppose E* ≡ *E*^∗^. *Then*, 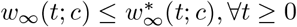.

#### Theorem 8

*Let T <* ∞. *Suppose E* ≡ *E*^∗^. *Then*, ∃ Δ *>* 0 *such that* 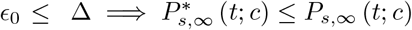 *for any t* ∈ [0, *T*]. *Moreover, we can estimate*

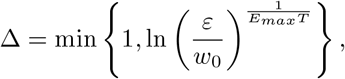

*where ε is a chosen bound on the maximal wild-type density difference*.

#### Theorem 9

*For any K <* ∞ *and t* ≥ 0, *we have that*

- 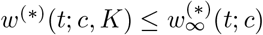
- 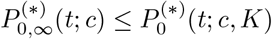
- 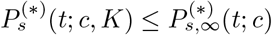
- 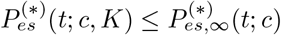

The proofs of Theorems 7, 8, and 9 are similar to those of Theorems 1, 2, and 3, respectively. Thus, we omit them to avoid redundancy.

## Footnotes

1 We show in Section SI 1 that the corresponding limit actually exists for our relevant parameter values.

2 Notations with superscript ^∗^ corresponds to the biostatic case

3 We slightly relax this condition in Theorem 4.

4 The derivations of these formulas are presented in SI 1

5 Although both AMR sources can be considered dose-dependent, we only require a bound on the *de novo* mutation rate.

6 Resistance that increases *rMIC* to a finite value higher than *zMIC* [1]

