## Supplementary Information for "Probability of Antibiotic Resistance During Treatment in Stochastic PK/PD-Based Bacterial Model with Distinct Drug and Mutation Modes"

Chimezie Izuazu     Cameron Browne*

June 3, 2026

The supplementary information contains the following:

1. **Establishment Probability:** The derivation of the establishment probability of the resistant bacterial strain.

2. **Effect of Partial Resistance:** Simulations considering partial resistance.

3. **Effect of De novo Mutation:** Simulations considering various HGT rates.

4. **PD–PK-Based Treatment for Longer Duration:** Simulations of PD–PK-based survival probabilities considering longer treatment duration.

5. **Effect of HGT:** Simulations considering various *de novo* mutation rates.

6. **Sensitivity Analysis:** Simulations of sensitivity of survival probability to various dose-dependent mutation parameters.

#### 1 Establishment Probability

We simplify the following:

$$P_{es}\left(c\right) = 1 - \left[\frac{I_\infty}{1+I_\infty}\right]^{n_0} e^{-\left[\int_{t_0}^{\infty} \frac{\epsilon_0 \alpha_w (K - w(\tau;c))w(\tau;c)}{K(1+I_\infty - I_\tau)} d\tau\right]}. \tag{1}$$

$$P^*_{es}\left(c\right) = 1 - \left[\frac{I^*_\infty}{1+I^*_\infty}\right]^{n_0} e^{-\left[\int_{t_0}^{\infty} \frac{\epsilon_0 \alpha_w \left(K(1-\frac{E^*}{\alpha_w}) - w^*(\tau;c)\right) w^*(\tau;c)}{K\left(1+I^*_\infty - I^*_\tau\right)} d\tau\right]}. \tag{2}$$

*

For the biocidal case, we have

$$\begin{aligned}
I_T &= \int_0^T \left[\mu_m e^{\int_0^t (\mu_m - \lambda_h(\tau))d\tau}\right] dt \\
&= \mu_m \int_0^T \left[e^{\int_0^t \left(\mu_m - \gamma_h \frac{w(\tau)}{K} - \lambda_m(\tau)\right)d\tau}\right] dt \\
&= \mu_m \int_0^T \left[e^{\int_0^t \left[\mu_m - \alpha_m + \left(\frac{\alpha_m - \gamma_h}{K}\right) w(\tau)\right]d\tau}\right] dt \\
&= \mu_m \int_0^T \left[e^{\left[(\mu_m - \alpha_m)t + \left(\frac{\alpha_m - \gamma_h}{K}\right)\int_0^t w(\tau)d\tau\right]}\right] dt
\end{aligned}$$

Moreover,

$$\begin{aligned}
\int_0^t w(\tau)d\tau &= \int_0^t \frac{w_0 K\rho(c)e^{\rho(c)\tau}}{K\rho(c) + w_0\alpha_w(1-\epsilon_0)(e^{\rho(c)\tau}-1)} d\tau \\
&= \frac{K}{\alpha_w(1-\epsilon_0)} \ln\left(\frac{K\rho(c) + w_0\alpha_w(1-\epsilon_0)(e^{\rho(c)t}-1)}{K\rho(c)}\right)
\end{aligned}$$

Thus,

$$I_T = \mu_m \int_0^T \left[\frac{K\rho(c) + w_0\alpha_w(1-\epsilon_0)(e^{\rho(c)t}-1)}{K\rho(c)}\right]^{\frac{\alpha_m-\gamma_h}{\alpha_w(1-\epsilon_0)}} \left[e^{(\mu_m-\alpha_m)t}\right] dt$$

In general, the integral does not simplify to elementary functions, but it can be written in terms of a hypergeometric function. Under the simplifying assumption that $\alpha_m = \alpha_w(1-\epsilon_0) + \gamma_h$, we have

$$\frac{I_\infty}{1+I_\infty} = \frac{\mu_m w_0\alpha_w(1-\epsilon_0) - \mu_m K[\varrho(c) + (\mu_m - \alpha_m)]}{\mu_m w_0\alpha_w(1-\epsilon_0) - \alpha_m K[\varrho(c) + (\mu_m - \alpha_m)]}$$

Therefore,

$$P_{es}(c) = 1 - \left[\frac{\mu_m w_0\alpha_w(1-\epsilon_0) - \mu_m K[\varrho(c) + (\mu_m - \alpha_m)]}{\mu_m w_0\alpha_w(1-\epsilon_0) - \alpha_m K[\varrho(c) + (\mu_m - \alpha_m)]}\right]^{n_0} e^{-\left[\int_{t_0}^{\infty} \frac{\epsilon_0\alpha_w(K-w(\tau;c))w(\tau;c)}{K(1+I_\infty - I_\tau)}d\tau\right]}.$$

Similarly, for the biostatic case, we have

$$P^*_{es}(c) = 1 - \left[\frac{\mu_m w_0\alpha_w(1-\epsilon_0) - \mu_m K[\varrho^*(c) + (\mu_m - \alpha_m)]}{\mu_m w_0\alpha_w(1-\epsilon_0) - \alpha_m K[\varrho^*(c) + (\mu_m - \alpha_m)]}\right]^{n_0} e^{-\left[\int_{t_0}^{\infty} \frac{\epsilon_0\alpha_w\left(K(1-\frac{E^*(c)}{\alpha_w}) - w^*(\tau;c)\right)w^*(\tau;c)}{K(1+I^*_\infty - I^*_\tau)}d\tau\right]}.$$

Further, $D(T;c) := \int_0^T \frac{\epsilon_0\alpha_w(K-w(\tau;c))w(\tau;c)}{K(1+I_\infty-I_\tau)}d\tau$ is an increasing function of $T$. Thus, $D(c) := \lim_{T\longrightarrow\infty} D(T;c) \in [0,\infty]$. In fact, we now establish that $D(c) \in [0,\infty)$ for the relevant parameter values.

$$I_T = \mu_m\left[\left(1 - \frac{w_0\alpha_w(1-\epsilon_0)}{K\varrho(c)}\right)\left(\frac{e^{(\mu_m-\alpha_m)T}-1}{(\mu_m-\alpha_m)}\right) + \frac{w_0\alpha_w(1-\epsilon_0)}{K\varrho(c)}\left(\frac{e^{[\varrho(c)+(\mu_m-\alpha_m)]T}-1}{[\varrho(c)+(\mu_m-\alpha_m)]}\right)\right]$$

$$I_\infty = \mu_m \left[ \frac{w_0 \alpha_w (1-\epsilon_0) - K\varrho(c)}{K\varrho(c)(\mu_m - \alpha_m)} - \frac{w_0 \alpha_w (1-\epsilon_0)}{K\varrho(c)\left[\varrho(c) + (\mu_m - \alpha_m)\right]} \right]$$

Thus,

$$I_\infty - I_T = \mu_m \left[ \frac{w_0 \alpha_w (1-\epsilon_0) - K\varrho(c)}{K\varrho(c)(\mu_m - \alpha_m)} e^{(\mu_m - \alpha_m)T} - \frac{w_0 \alpha_w (1-\epsilon_0)}{K\varrho(c)\left[\varrho(c) + (\mu_m - \alpha_m)\right]} e^{[\varrho(c) + (\mu_m - \alpha_m)]T} \right]$$

For simplicity, we set $A = \frac{\mu_m (w_0 \alpha_w (1-\epsilon_0) - K\varrho(c))}{K\varrho(c)(\mu_m - \alpha_m)}$, $a = (\mu_m - \alpha_m)$, $B = \frac{\mu_m w_0 \alpha_w (1-\epsilon_0)}{K\varrho(c)[\varrho(c) + (\mu_m - \alpha_m)]}$, and $b = [\varrho(c) + (\mu_m - \alpha_m)]$. Thus,

$$1 + I_\infty - I_T = 1 + Ae^{aT} - Be^{bT}, \text{with } a < 0, b < 0, A > 0, 1 > B > 0.$$

$$\begin{aligned}
\frac{\epsilon_0 \alpha_w (K - w(\tau;c)) w(\tau;c)}{K(1 + I_\infty - I_\tau)} &= \frac{\epsilon_0 \alpha_w (K - w(\tau;c)) w(\tau;c)}{K(1 + Ae^{a\tau} - Be^{b\tau})} \\
&\leq \frac{\epsilon_0 \alpha_w K w(\tau;c)}{K(1 + Ae^{a\tau} - Be^{b\tau})} \\
&= \frac{\epsilon_0 \alpha_w w(\tau;c)}{(1 + Ae^{a\tau} - Be^{b\tau})} \\
&\leq \frac{\epsilon_0 \alpha_w w_0}{(1 - Be^{b\tau})}.
\end{aligned}$$

$$\begin{aligned}
\int_0^T \frac{\epsilon_0 \alpha_w (K - w(\tau;c)) w(\tau;c)}{K(1 + I_\infty - I_\tau)} d\tau &\leq \int_0^\infty \frac{\epsilon_0 \alpha_w (K - w(\tau;c)) w(\tau;c)}{K(1 + I_\infty - I_\tau)} d\tau, \forall\, T \geq 0 \\
&\leq \int_0^\infty \frac{\epsilon_0 \alpha_w w_0}{(1 - Be^{b\tau})} d\tau. \\
&= \lim_{T \longrightarrow \infty} \int_0^T \frac{\epsilon_0 \alpha_w w_0}{(1 - Be^{b\tau})} d\tau. \\
&= \epsilon_0 \alpha_w w_0 \left[ \lim_{T \longrightarrow \infty} \int_0^T \frac{1}{(1 - Be^{b\tau})} d\tau \right]. \\
&= \epsilon_0 \alpha_w w_0 \left[ \lim_{T \longrightarrow \infty} \left( \frac{1}{b} \ln \left( \frac{-Be^{b\tau}}{1 - Be^{b\tau}} \right) \Bigg|_0^T \right) \right]. \\
&\leq \epsilon_0 \alpha_w w_0 \left[ \lim_{T \longrightarrow \infty} \left( \frac{1}{b} \ln \left( \frac{1 - B}{1 - Be^{bT}} \right) \right) \right]. \\
&\leq \epsilon_0 \alpha_w w_0 \left[ \lim_{T \longrightarrow \infty} \left( \frac{1}{-b} \ln \left( \frac{1 - Be^{bT}}{1 - B} \right) \right) \right]. \\
&= \frac{\epsilon_0 \alpha_w w_0}{-b} \ln \left( \frac{1}{1 - B} \right). \\
&= \ln \left( \frac{1}{1 - B} \right)^{\frac{\epsilon_0 \alpha_w w_0}{-b}}.
\end{aligned}$$

Thus, by the monotone convergence theorem, $D(c) \in [0, \infty)$. Similarly, for the biostatic treatment, we obtain that $D^*(c) \in [0, \infty)$.

### 2 Effect of Partial Resistance

In the main text, we have assumed the worst possible case, i.e., total antibiotic resistance; however, a more realistic setting should consider partial antibiotic resistance (i.e., resistance that increases $rMIC$ to a finite value compared with $zMIC$ [1]). Consequently, we present some numerical simulations considering various partial resistance levels and sources.

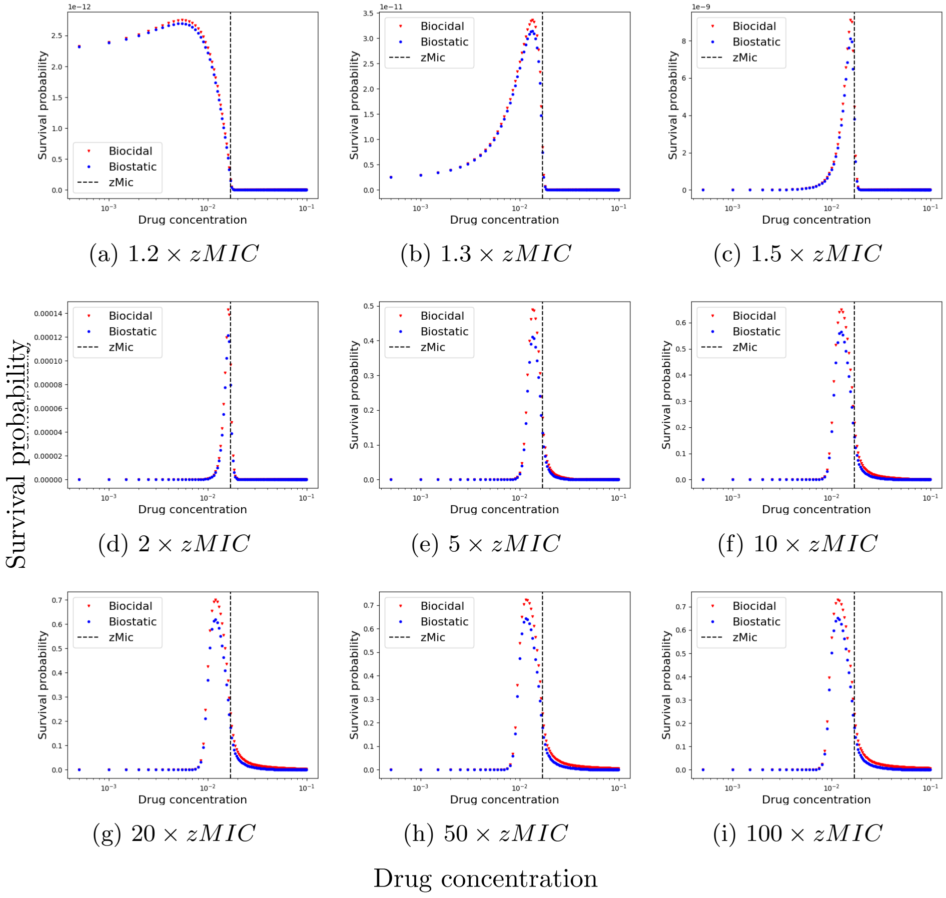


(a) $1.2 \times zMIC$ (b) $1.3 \times zMIC$ (c) $1.5 \times zMIC$

(d) $2 \times zMIC$ (e) $5 \times zMIC$ (f) $10 \times zMIC$

(g) $20 \times zMIC$ (h) $50 \times zMIC$ (i) $100 \times zMIC$

Figure 1: Survival probability with respect to drug concentration for 72-h PD biostatic vs. biocidal treatments considering only partial *de novo* resistance (i.e., no HGT and SGV).

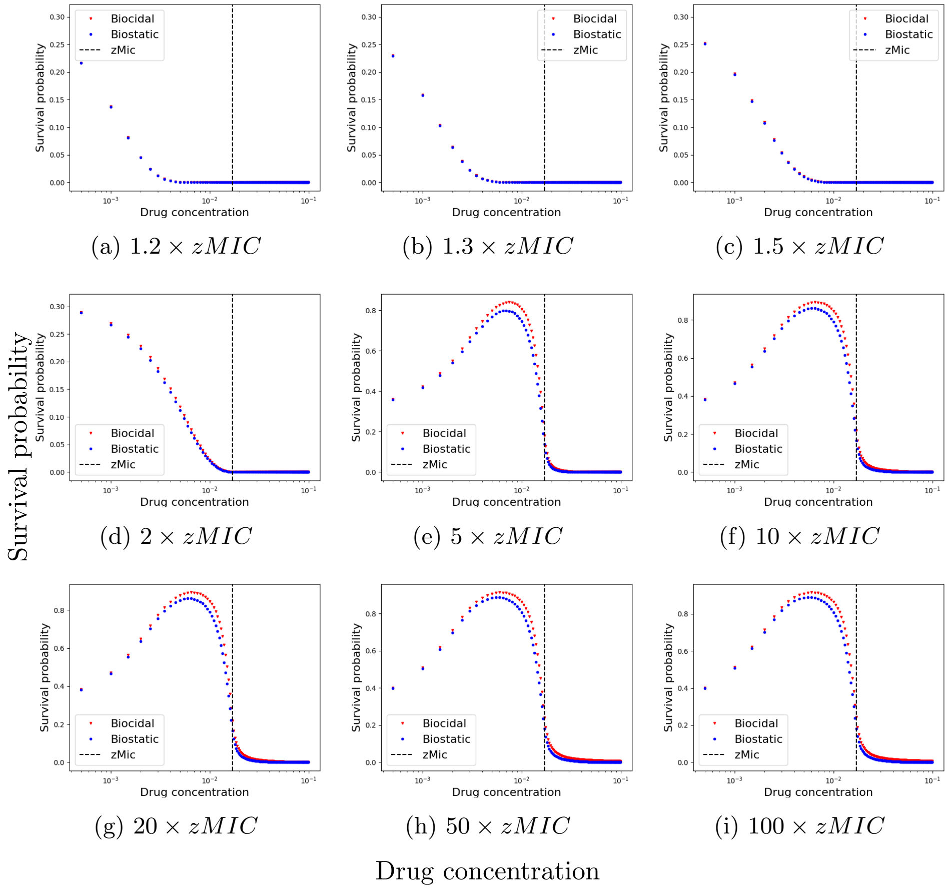


Figure 2: Survival probability with respect to drug concentration for 72-h PD biostatic vs. biocidal treatments considering partial *de novo* resistance and HGT ($\gamma_h = 1.5$).

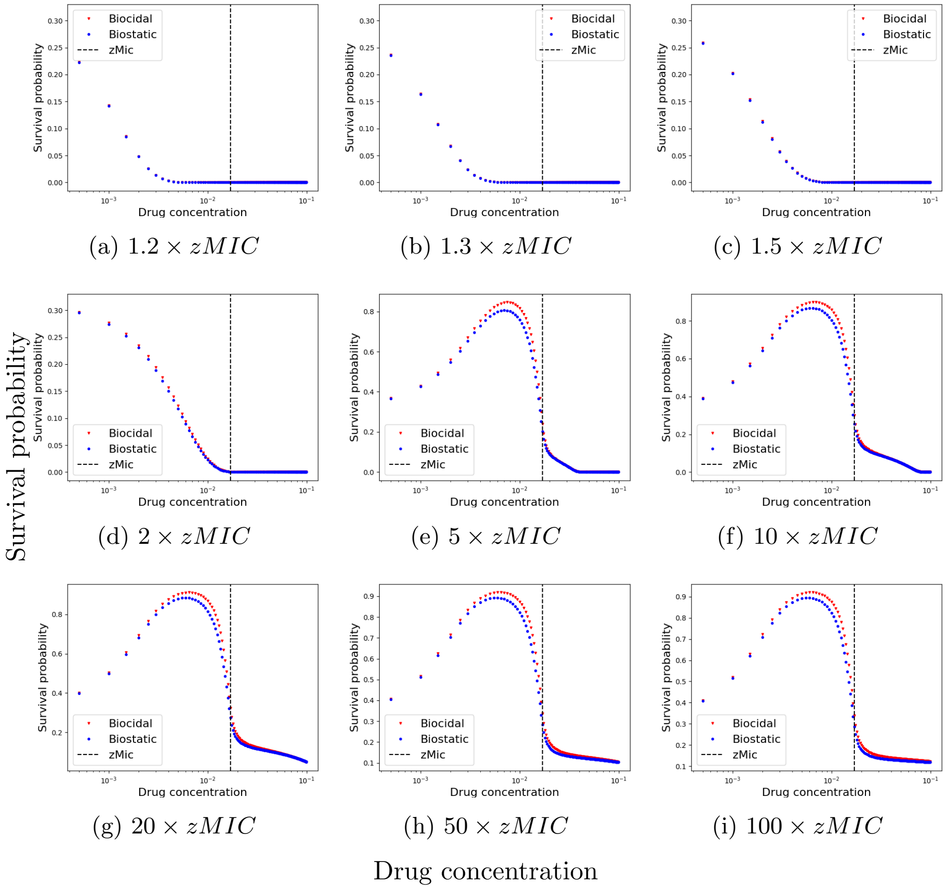


(a) $1.2 \times zMIC$ (b) $1.3 \times zMIC$ (c) $1.5 \times zMIC$

(d) $2 \times zMIC$ (e) $5 \times zMIC$ (f) $10 \times zMIC$

(g) $20 \times zMIC$ (h) $50 \times zMIC$ (i) $100 \times zMIC$

Figure 3: Survival probability with respect to drug concentration for 72-h PD biostatic vs. biocidal treatments considering partial *de novo* resistance, HGT ($\gamma_h = 1.5$), and SGV ($n_0 = 1$).

### 3 Effect of De novo Mutation

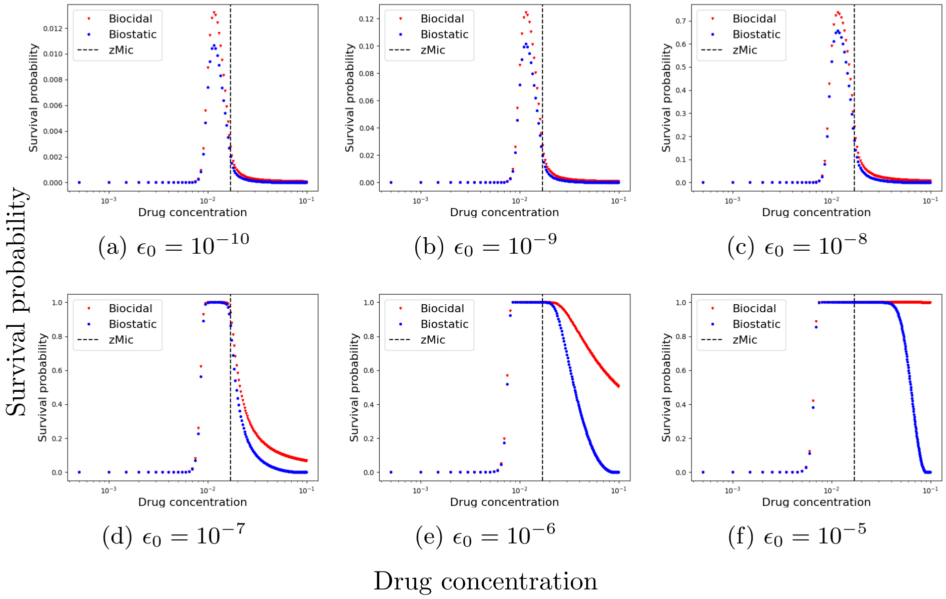


(a) $\epsilon_0 = 10^{-10}$ (b) $\epsilon_0 = 10^{-9}$ (c) $\epsilon_0 = 10^{-8}$

(d) $\epsilon_0 = 10^{-7}$ (e) $\epsilon_0 = 10^{-6}$ (f) $\epsilon_0 = 10^{-5}$

Figure 4: Survival probability with respect to drug concentration for 72-h PD biostatic vs. biocidal treatments considering various *de novo* resistance rates.

### 4 PK/PD-Based Treatment for Longer Duration

Survival probability

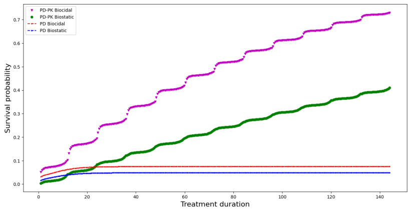


(a) *De novo* survival chance

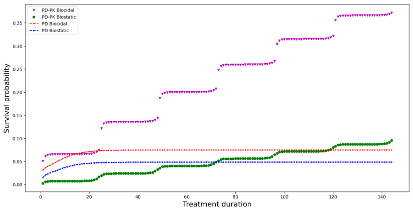


(b) *De novo* survival chance

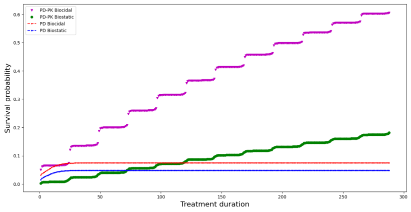


(c) *De novo* survival chance

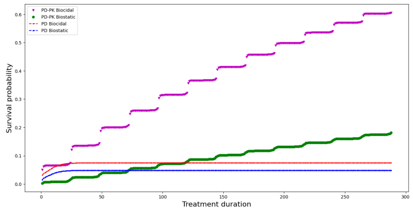


(d) *De novo* survival chance

Treatment duration

Figure 5: Comparison of 12-h (first column) and 24-h (second column) periodic dosing with $c_0 = 0.062$ and a corresponding constant concentration that matches the AUCTC.

### 5 Effect of HGT

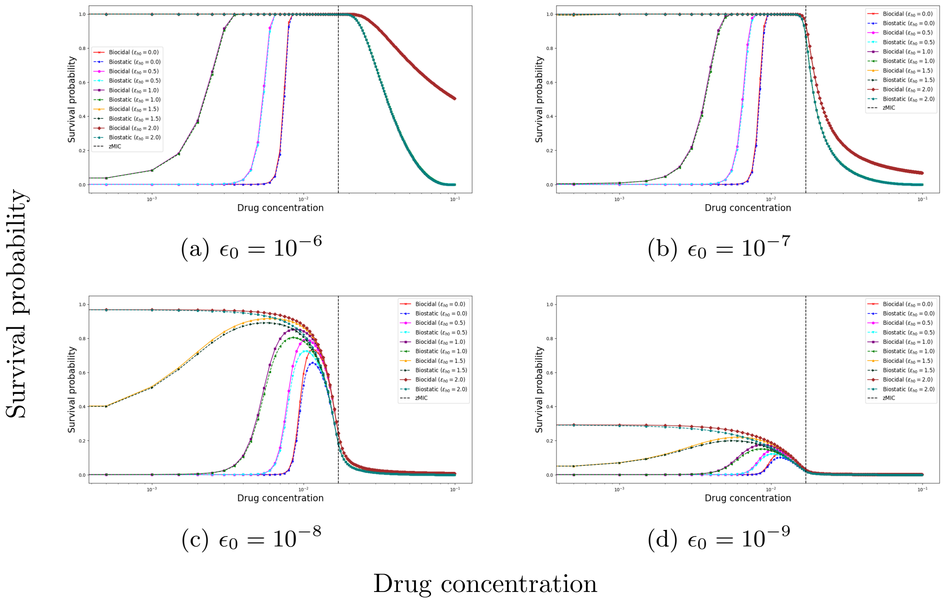


Figure 6: Effect of HGT on survival probability with respect to drug concentration for 72-h PD biostatic vs. biocidal treatments considering various *de novo* resistance rates.

Figure (6) indicates that the impact of HGT on survival probability depends strongly on the baseline *de novo* resistance rate. Although HGT has little influence when resistance emerges frequently, its effect becomes increasingly pronounced as the baseline resistance rate decreases, indicating that HGT can serve as a critical alternative pathway for resistance emergence.

### 6 Sensitivity Analysis

| Parameter | Meaning | Value (unit) | Source | Considered range |
|---|---|---|---|---|
| $\epsilon^{max}$ | Maximum mutation rate for nonlinear mutation | $10^{-4.06}$ | [2] | $[10^{-6}, 10^{-4}]$ |
| $\kappa_1$ | Hill coefficient | 2.34 | [2] | $[1,\ 2.5]$ |
| $c_2^{max}$ | Drug concentration to attain half of $\epsilon^{max}$ | 50 $\mu gml^{-1}$ | [2] | $[0.005,\ 60]$ |
| $\epsilon_0$ | Intercept for linear mutation | $2.15 \times 10^{-10}, 3.02 \times 10^{-8}$ | [3] | $[10^{-11}, 10^{-7}]$ |
| $\theta$ | Slope for linear mutation | $7.46 \times 10^{-12}, 1.86 \times 10^{-10}$ | [3] | $[10^{-13}, 10^{-9}]$ |

Table 1: Dose-dependent mutation parameters

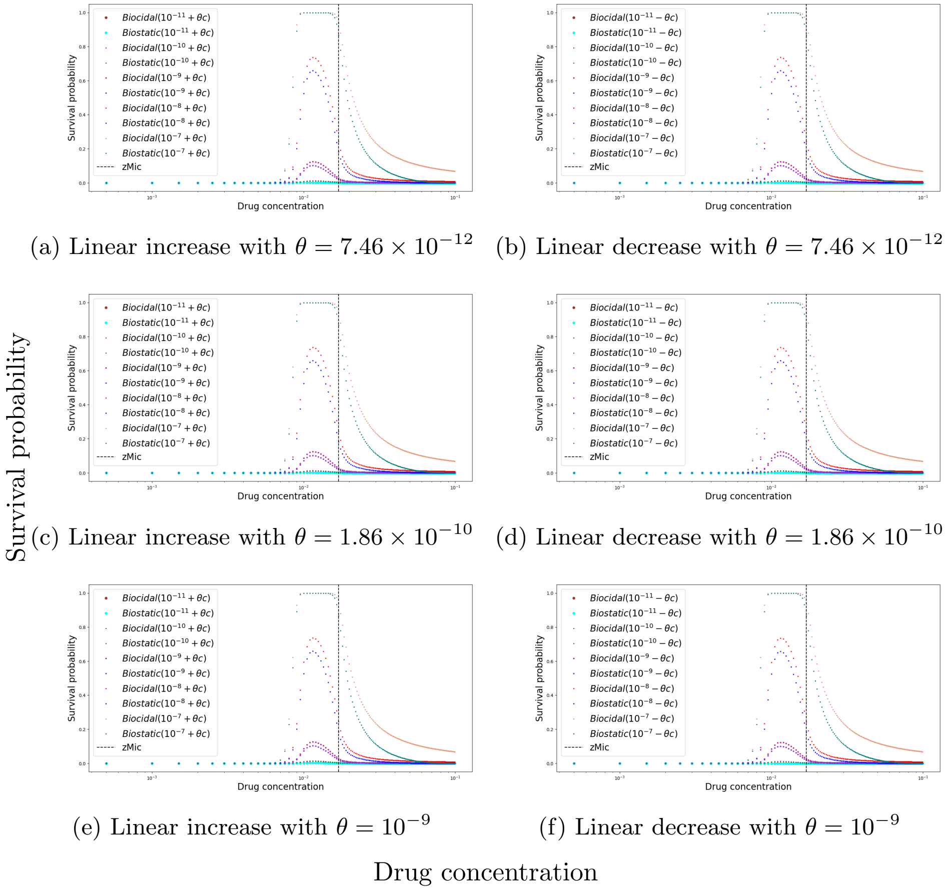


(a) Linear increase with $\theta = 7.46 \times 10^{-12}$ (b) Linear decrease with $\theta = 7.46 \times 10^{-12}$

(c) Linear increase with $\theta = 1.86 \times 10^{-10}$ (d) Linear decrease with $\theta = 1.86 \times 10^{-10}$

(e) Linear increase with $\theta = 10^{-9}$ (f) Linear decrease with $\theta = 10^{-9}$

Figure 7: Survival probability with respect to drug concentration for 72-h PD biostatic vs. biocidal treatments under various linear dose-dependent mutation parameter settings.

Figure 7 shows the survival probabilities under the linear dose-dependent mutation rate considering various parameters adapted from the literature [3]. Remarkably, there are no notable change in survival probability for various slopes of the linear decreasing and increasing mutation rates. Intuitively, the magnitude of the "intermediate-dose peak" increases with the initial mutation rate (intercept).

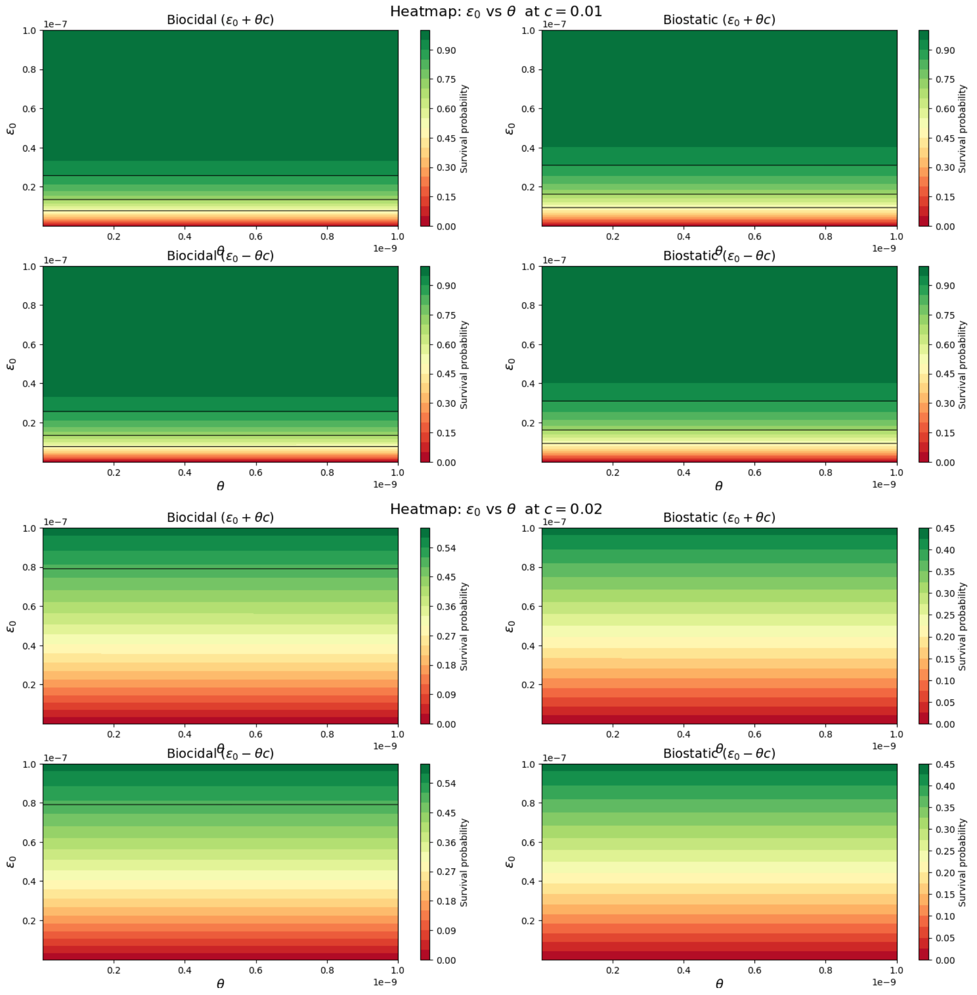


Figure 8: Heatmap for sensitivity of survival probability to linear dose-dependent mutation parameters for 72-h PD biostatic vs. biocidal treatments.

Figure (8) shows that the survival probability is primarily driven by the slope of the linear dose-dependent mutation function, with comparatively weak sensitivity to the intercept. This trend is consistent across both biostatic and biocidal treatments and for the parameter sets considered.

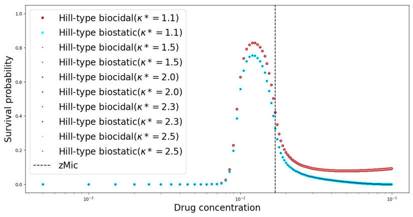


(a) Nonlinear ($\epsilon^{max} = 10^{-4.06}; c_2^{max} = 50$)

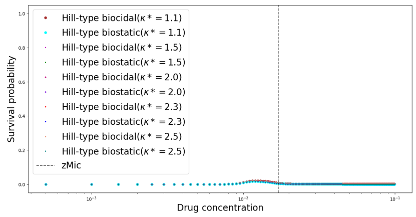


(b) Nonlinear ($\epsilon^{max} = 10^{-6}; c_2^{max} = 50$)

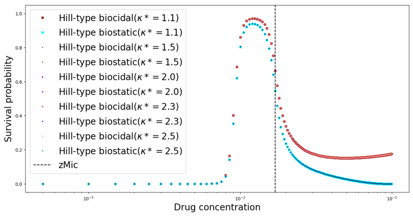


(c) Nonlinear ($\epsilon^{max} = 10^{-4.06}; c_2^{max} = 25$)

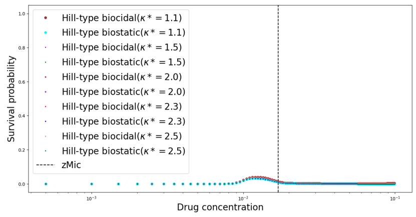


(d) Nonlinear ($\epsilon^{max} = 10^{-6}; c_2^{max} = 25$)

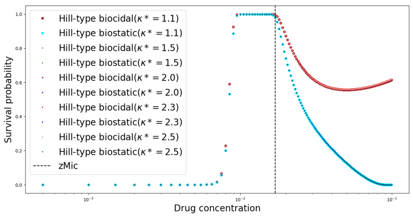


(e) Nonlinear ($\epsilon^{max} = 10^{-4.06}; c_2^{max} = 5$)

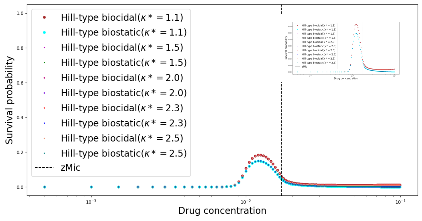


(f) Nonlinear ($\epsilon^{max} = 10^{-6}; c_2^{max} = 5$)

Survival probability

Drug concentration

Figure 9: Survival probability with respect to drug concentration for 72-h PD biostatic vs. biocidal treatments under various nonlinear dose-dependent mutation parameter settings.

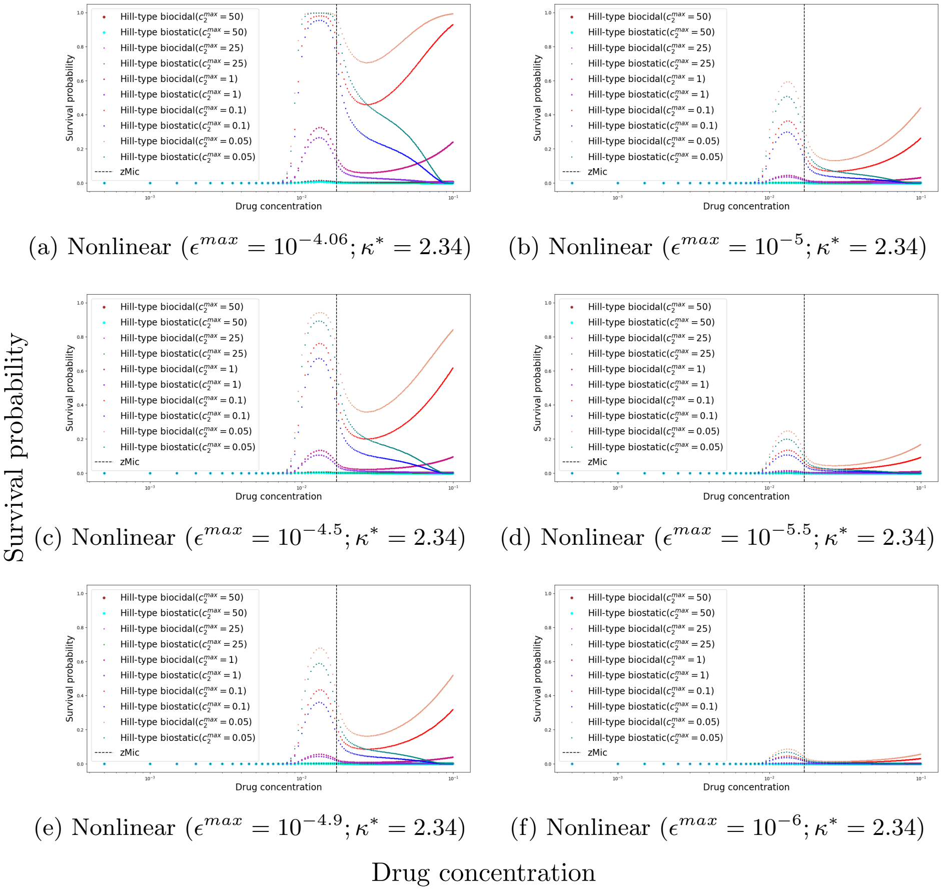


(a) Nonlinear ($\epsilon^{max} = 10^{-4.06}; \kappa^* = 2.34$) (b) Nonlinear ($\epsilon^{max} = 10^{-5}; \kappa^* = 2.34$)

(c) Nonlinear ($\epsilon^{max} = 10^{-4.5}; \kappa^* = 2.34$) (d) Nonlinear ($\epsilon^{max} = 10^{-5.5}; \kappa^* = 2.34$)

(e) Nonlinear ($\epsilon^{max} = 10^{-4.9}; \kappa^* = 2.34$) (f) Nonlinear ($\epsilon^{max} = 10^{-6}; \kappa^* = 2.34$)

Drug concentration

Figure 10: Survival probability with respect to drug concentration for 72-h PD biostatic vs. biocidal treatments under various nonlinear dose-dependent mutation parameter settings.

Figures 9 and 10 show the survival probabilities under the nonlinear dose-dependent mutation rate considering various parameters adapted from the literature [2]. Without HGT (See main text figures), the figures show that the survival probability is insensitive to the Hill coefficient, $\kappa^*$. Notably, the magnitude of the "intermediate-dose peak" decreases with an increase in the drug concentration to reach half of the maximum mutation rate, $c_2^{max}$. Interestingly, at lower $c_2^{max}$ values, the survival probability for the biocidal treatment tends to increase at supra-MIC values; the rate of increase is accelerated with a decrease in $c_2^{max}$. This phenomenon supports our claim that the biostatic treatment is more effective in resistance suppression. Figure (11) shows the sensitivity of the survival probability to the parameters of the nonlinear dose-dependent mutation function. Notably, the first two sub-figures exhibit nearly uniform color

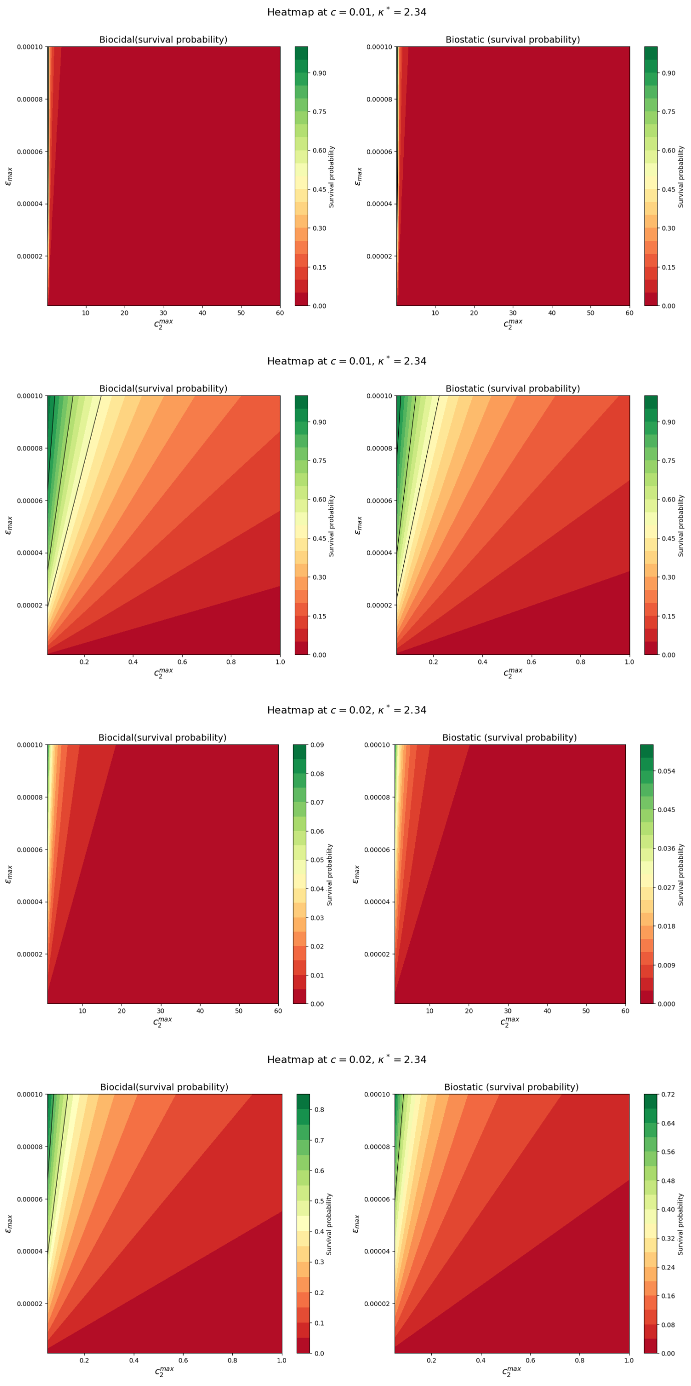


Figure 11: Heatmap for sensitivity of survival probability to nonlinear dose-dependent mutation parameters for 72-h PD biostatic vs. biocidal treatments.

distributions, indicating that the survival probability is relatively insensitive to parameter variations in this regime. Meanwhile, the remaining sub-figures display pronounced diagonal gradients, suggesting that the survival probability depends strongly on the combined effects of both nonlinear mutation parameters. In these cases, increasing either parameter generally results in a higher survival probability. Notably, similar qualitative trends are observed for both biostatic and biocidal treatments. Unlike the linear case (Figure (8)), where the slope parameter primarily drives the response, the nonlinear formulation exhibits a stronger interaction between parameters, producing the diagonal sensitivity patterns observed across both biostatic and biocidal treatments.
